# Functional Muscle Networks as Biomarkers of Post-Stroke Motor Impairment and Therapeutic Responsiveness

**DOI:** 10.1101/2025.08.17.670726

**Authors:** David O’Reilly, Giorgia Pregnolato, Andrea Turolla, Pawel Kiper, Ioannis Delis, Giacomo Severini

## Abstract

Standardised assessment of post-stroke motor impairment and treatment responsiveness remains a major clinical challenge. In this study, we tackle this challenge by applying a novel muscle network analysis framework to stroke survivors undergoing intensive upper-limb motor training. Our approach revealed distinct patterns of redundant and synergistic muscle interactions, collectively reflecting the diverse biomechanical roles of flexor- and extensor-driven networks. From these patterns, we derived new biomarkers that stratified patients by gross motor impairment severity and therapeutic responsiveness, each associated with unique physiological signatures. Remarkably, we identified a shift from redundancy to synergy in muscle coordination as a hallmark of effective motor recovery — a transformation supported by a more precise quantification of impairment over conventional approaches. These findings offer an in-depth characterisation of post-stroke motor recovery and establish a robust, independent tool for evaluating rehabilitation efficacy. Future research should employ this framework to identify biomarkers of activities- and participation-related functional recovery.

## Introduction

Stroke is a leading cause of death and disability worldwide, the incidence of which is predicted to rise globally in coming years (Saini et al., 2021). Stroke induces motor impairments among survivors, characterised by hemiparesis (i.e. unilateral loss of function especially of the upper limb), muscle weakness, dyscoordination and fatigue (Sathian et al., 2011; Teasell & Hussein, 2016). Clinical practice primarily focuses on early and intensive intervention to maximise movement restoration (Shiromoto et al., 2017; Teasell et al., 2012), however recent evidence advocates for the importance of long-term rehabilitation (Ballester et al., 2019; Kwakkel et al., 2023; Teasell et al., 2012). Nevertheless, current clinical assessment scales don’t effectively convey comprehensive information about stroke survivors at all recovery stages, partly due to the reliance of clinically important change metrics on acute-stage recovery rates, their inability to distinguish genuine recovery from compensation or to capture participation-level outcomes (Ballester et al., 2019; Bushnell et al., 2015; Page et al., 2012). Hence, a current research gap lies in effective assessment tools that effectively quantify post-stroke motor impairment and treatment responsiveness at all recovery stages.

Muscle synergy analysis is a data-driven and theoretically supported approach from the motor neurosciences that quantifies coordinated patterns of muscle activity (‘*muscle synergies*’) from surface electromyographic (sEMG) signals (Berret et al., 2019; Bruton & O’Dwyer, 2018; Cheung & Seki, 2021), interpreting these patterns as the output of functionally modular neural networks in the central nervous system. Through this approach, people affected by stroke have been shown to exhibit different neurophysiological responses depending on impairment severity and adaptations during recovery, including the preservation, merging and fractionation of unimpaired motor patterns along with the generation of novel motor patterns (Cheung et al., 2012; Clark et al., 2010; García-Cossio et al., 2014; Hashiguchi et al., 2016; Hong et al., 2021; Pregnolato et al., 2025). Indeed, useful biomarkers have been produced from this approach that convey linear relationships with clinically validated measures of motor impairment (Clark et al., 2010; Schwartz et al., 2016). However, restrictive model assumptions (e.g. linearity) and the lack of a direct mapping of muscle interactions to task performance has hampered the progression of this approach in practical applications (Alessandro et al., 2013; Berret et al., 2019; Cheung & Seki, 2021; de Rugy et al., 2013). In our recent work (O’Reilly & Delis, 2024; Ó’Reilly & Delis, 2022), we addressed these important limitations by incorporating network- and information-theoretic approaches into a novel, generalised framework (Network-Information Framework (NIF)) that maps muscle interactions directly to their functional consequences (Fig.1(A)). In doing so, we can more precisely quantify functional muscle co-variations, distinguishing between task-relevant (i.e. muscle activation patterns associated with task-specific movements (pink-orange chequerboard intersection (Fig.1(A))) from task-irrelevant (i.e. muscle activation patterns present across motor tasks (yellow intersection exclusively shared by *m*_*x*_ and *m*_*y*_ (Fig.1(A)) contributions intermixed in conventional approaches and going further in characterising their functional roles as functionally-similar (i.e. redundant (pink in pink-orange chequerboard intersection (Fig.1(A)) and -complementary (i.e. synergistic (orange in pink-orange chequerboard intersection (Fig.1(A))). Hence, the NIF offers novel research opportunities and practical use cases for muscle synergy analysis that may be beneficial in addressing these current limitations in the clinical assessment of the stroke population.

**Figure 1.**
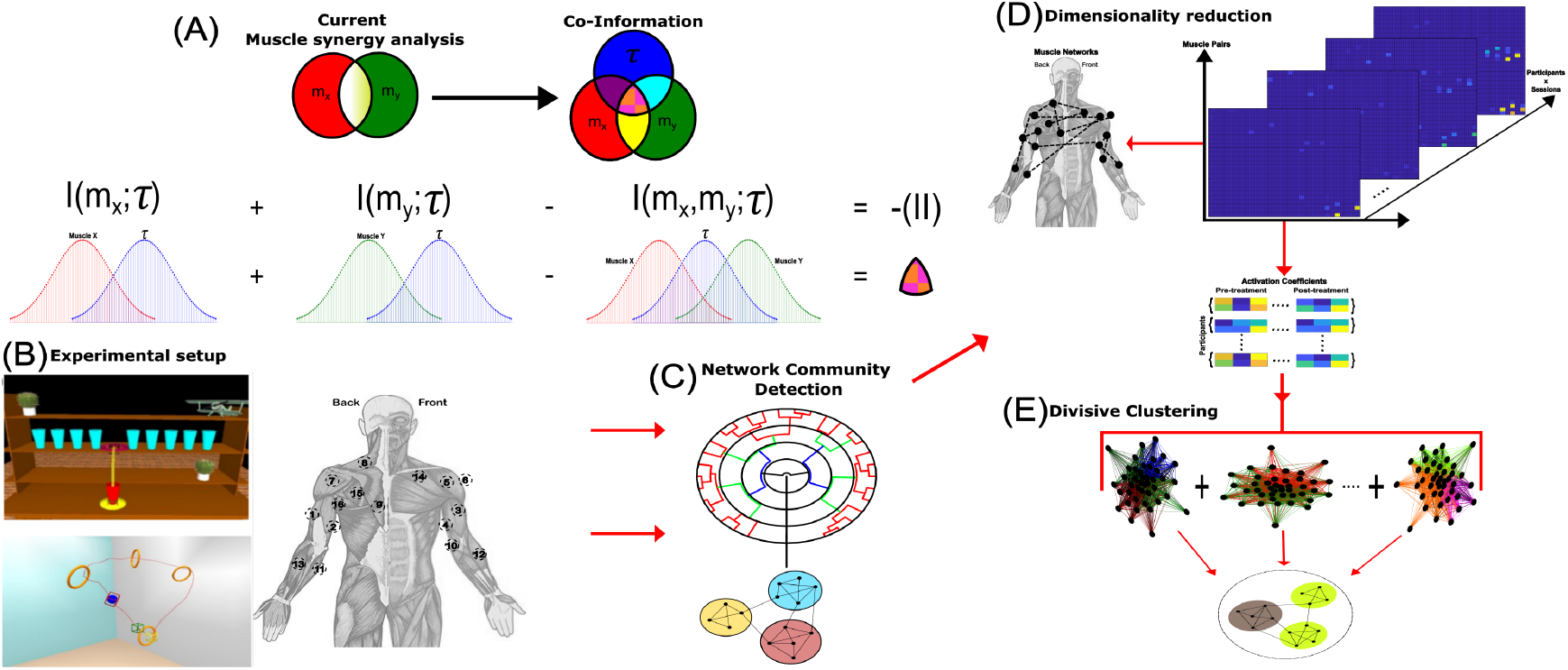
**(A)** We apply a novel muscle network analysis approach that leverages Co-Information (*II*) to map muscle pairs (*m*_*x*_, *m*_*y*_) to task performance (*τ*) (O’Reilly & Delis, 2024), thus dissecting the task-relevant information (pink-orange chequerboard intersection) between muscles from task-irrelevant information (yellow intersection shared exclusively by *m*_*x*_ and *m*_*y*_), and characterising their functional roles as either net functionally-similar (i.e. redundant) or -complementary (i.e. synergistic). **(A)** *II* characterises each muscle interaction as either net redundant (negative values) or synergistic (positive values) by contrasting the total mutual information each muscle shares with *τ* separately (*I*(*m*_*x*_; *τ*) + *I*(*m*_*y*_; *τ*)) against the task information observed when *m*_*x*_ and *m*_*y*_ are combined (*I*(*m*_*x*_, *m*_*y*_; *τ*)) (Ince et al., 2017; McGill, 1954). **(B)** An example virtual scenario from the virtual reality treatment where participants interacted with a real manipulandum to perform motor tasks such as placing a cup on a shelf. The correct movement trajectory (yellow line) was illustrated, which participants had to emulate. Task complexity was enhanced by including different objects and barriers, requiring participants to recruit different muscle groups to emulate the trajectory. Before and after the intervention, sEMGs were recorded from 16 muscles: triceps lateralis (**1**=TL), triceps medialis (**2**=TM); biceps short-head (**3**=BS), biceps long-head (**4**=BL); anterior deltoid (**5**=AD); lateral deltoid (**6**=LD); posterior deltoid (**7**=PD); upper trapezius (**8**=UT); rhomboid major (**9**=RM); brachialis (**10**=BRACH); supinator (**11**=SU); brachioradialis (**12**=BR); pronator teres (**13**=PT); pectoralis major (**14**=PM); infraspinatus (**15**=Infra); teres major (**16**=TEM) on both affected and unaffected sides, while participants performed 10 repetitions of seven standardised motor tasks (see ‘*Experimental setup and Data collection*’ in Materials and Methods) (Maistrello et al., 2021; Pregnolato et al., 2025). **(C)** *II* is quantified between all sEMG pairs to generate functional muscle networks, the modular structure of which is determined across participants and sessions using network community detection (Ahn et al., 2010). **(D)** The number of functional modules identified serves as the input parameter into a dimensionality reduction algorithm—projective non-negative matrix factorisation (PNMF)—to extract network components and their underlying session- and participant-specific activation coefficients (see ‘*Extraction of Redundant and Synergistic Muscle Networks*’ in Materials and Methods) (Ó’Reilly & Delis, 2022; Yang & Oja, 2010). **(E)** The activation coefficients are input into a novel divisive clustering algorithm to identify patient clusters both within (impairment clusters) and between (treatment response clusters) pre- and post-treatment (see ‘*Clustering stroke survivors based on impairment severity and therapeutic responsiveness*’ in Materials and Methods) (O’Reilly & Delis, 2025).

Therefore, in the current study we applied the NIF to a cohort of 42 stroke survivors performing 7 standardised upper-limb movements pre- and post-20 sessions of intensive upper-limb motor training through either a virtual reality (VR) or conventional physical therapy (PT) intervention (Fig.1(B)). In doing so, we aimed to provide novel characterisations and biomarkers of post-stroke impairment and therapeutic responsiveness that address these current limitations in post-stroke clinical assessments. To do so, here we focussed our analysis on quantifying task-dependent muscle couplings (collectively referred to as *II*), extracting functionally-similar (i.e. redundant) and -complementary (i.e. synergistic) modules and performing an in-depth analysis on their underlying activation patterns (Fig.1(A-E)). These functional modules reflected the diverse biomechanical contributions of collective muscle interactions towards 7 standardised motor tasks on both affected- and unaffected sides. Through our novel clustering algorithm (Fig.1(E)), we identified patient clusters aligned with and accentuating established clinical measures of gross motor impairment (moderately and severely-impaired subgroups) and treatment responsiveness (responders and non-responders subgroups). Finally, by probing the underlying contributions of merging, fractionation, and preservation patterns to the observed patient clusters, we shed mechanistic light on their role in the manifestation of movement pathology. Taken together, this study provides an in-depth functional characterisation of post-stroke impairment and treatment responsiveness whilst progressing the clinical use case for muscle synergy analysis as an independent assessment tool for post-stroke rehabilitation.

## Results

The cohort of stroke survivors overall experienced a statistically significant increase at FMA-UE (Pre-treatment: 43.1±13.2, Post-treatment: 49.1±13.6 (t= -7.84, *p* < 0.001)), representing a clinically important effect from rehabilitation on the gross motor functions of the upper-extremity (Page et al., 2012). Although the VR group displayed a greater improvement in FMA-UE scores compared to PT, no significant differences between treatment conditions were found at baseline (PT: 42.75±16.3, VR: 43.2±12.6 (t= -0.69, *p* > 0.9)) or at follow-up (PT: 45.62±15.2, VR: 49.9±13.3 (t= -0.723, *p* > 0.45)). 21 responders and 21 non-responders were identified using the conventional MCID of >5 points on the FMA-UE (responders (PT=3/8, VR=18/34), non-responders (PT=5/8, VR=16/34)) (Page et al., 2012). No significant differences were found between VR and PT responders and non-responders at baseline or follow-up (*p* > 0.05).

### Functional muscle networks reflect the biomechanical contributions of collective muscle interactions

The application of the NIF across the post-stroke cohort and both pre- and post-sessions identified five redundant (R1-R5 (Supplementary Materials Fig.1)) and seven synergistic (S1-S7 (Supplementary Materials Fig.2)) networks of co-occurring muscle interactions across affected and unaffected sides. These functional muscle networks were comprised of muscle interactions that contributed in functionally-similar and -complementary ways respectively towards the seven standardised motor tasks performed (see Table.2 of Materials and Methods section for details). Subsequent examination of the muscle interaction strengths (relative network edge width), principal muscles (relative node size) and submodular structure (node colour) enabled a biomechanical function to be attributed to each extracted component (see the supporting information for Supplementary Materials Fig.1-2 for detailed descriptions and ‘*Extraction of Redundant and Synergistic Muscle Networks*’ section of the Materials and Methods section for technical details). Muscle acronyms are described in the caption of Fig.1 here.

To briefly summarise, the post-stroke cohort were characterised by multiple extensor- and flexor-driven muscle networks comprised of predominantly redundant and synergistic muscle interactions respectively. The redundant muscle networks specifically were also consistently composed of functional contributions of shoulder girdle and/or elbow stabilisers (e.g. TEM, Infra, PT and SU) supporting prime-mover musculature (e.g. AD, TL and TM). There was a functional correspondence between the affected- and unaffected-side redundant networks (e.g. R2 and R5 (Supplementary Materials Fig.1)), however, noticeable differences relating to the underlying impairment of the affected-side across the cohort were also represented (e.g. R3 (Supplementary Materials Fig.1), involving interactions between SU and the shoulder musculature on both sides, displayed a specific reliance on PM on the affected-side whereas for the unaffected-side, the interactions were spread across PM, AD and LD, representing the characteristic loss of muscle selectivity at the shoulder level post-stroke). Meanwhile the synergistic muscle networks were generally characterised by widespread dependencies between distal and proximal musculature (e.g. S6 (Affected-side) Supplementary Materials Fig.2) that were less comparable across affected- and unaffected-sides. Contributions from joint stabilising musculature such as TEM and Infra were also found among synergistic networks, suggesting they contribute in both functionally-similar and -complementary ways to task performance with respect to prime-mover muscles.

### Network activation patterns are associated with gross motor impairment, treatment type and responsiveness

In assessing the statistical relationships between the activation of specific redundant and synergistic network components and gross motor functional impairment, we found that the degree of activation of S2 and S6 (both comprised of complementary muscular contributions to elbow flexion with shoulder girdle stabilisation (Fig.2(A)) and Supplementary Materials Fig.2) at baseline was associated with lower FMA-UE scores both at baseline (*β* = -174.5±62.5 (*p* = 0.008), *β* = -159.7±45.6 (*p* = 0.001) respectively) and at follow-up (*β* = -172.5±65.4 (*p* = 0.012), *β* = -155.1±48.2 (*p* = 0.003) respectively) (Fig.2(A)). This suggests that specific networks of synergistic muscle interactions are strongly linearly related to impairment at the group-level in such a way that persists after clinical intervention. Meanwhile, no significant relationships were found between individual activation coefficients and FMA-UE scores after treatment, suggesting that perhaps the activation coefficients after treatment reflected features specific of participant and modality provided.

**Figure 2.**
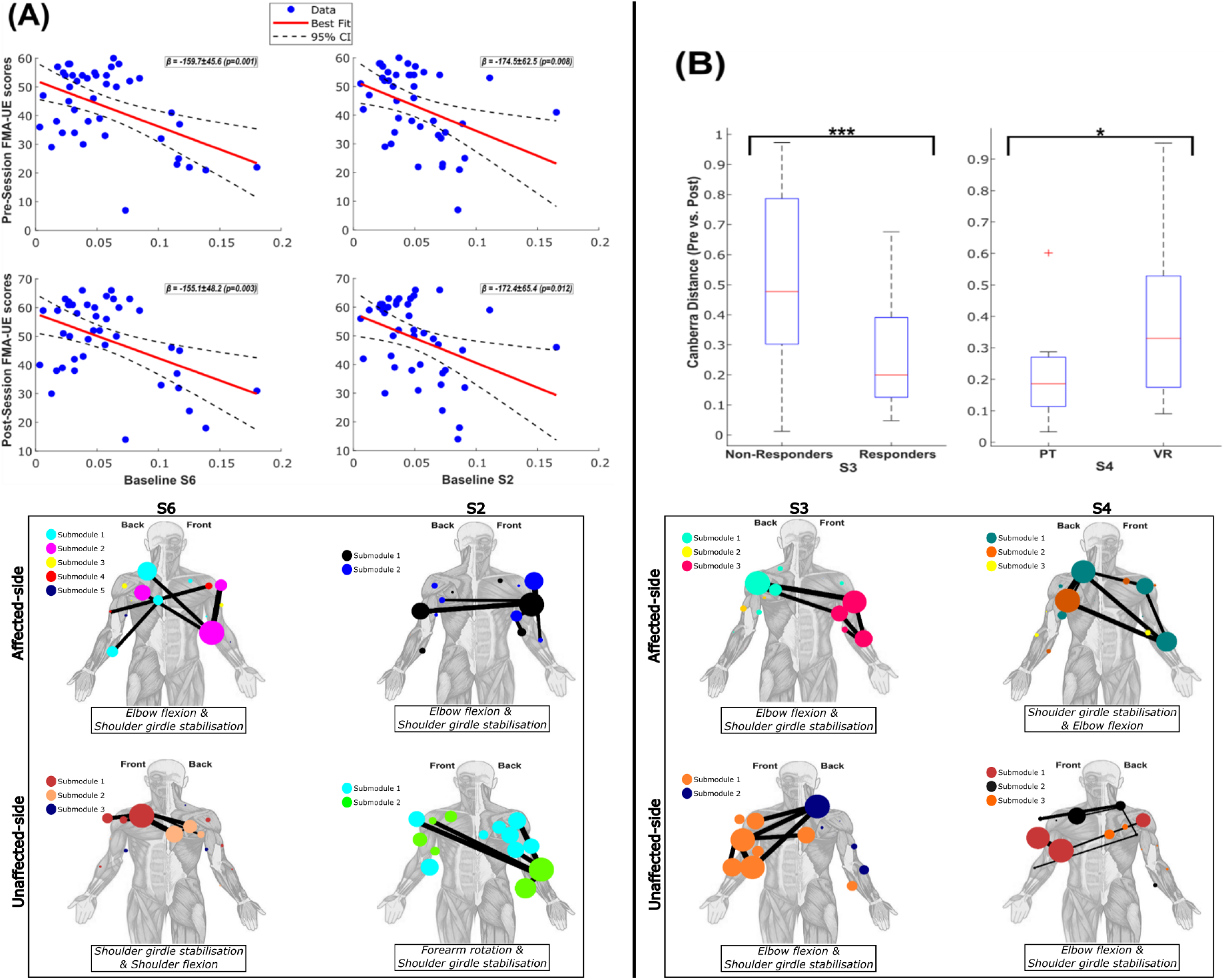
**(A)** Univariate linear regression analyses revealed the activation of both S2 and S6 at baseline was significantly associated with lower FMA-UE scores at baseline (*β* = -174.5±62.5 (*p* = 0.008), *β* = -159.7±45.6 (*p* = 0.001) respectively) and follow-up (*β* = -172.5±65.4 (*p* = 0.012), *β* = -155.1±48.2 (*p* = 0.003) respectively). No significant relationships were found between activation patterns and FMA-UE scores after stroke. The affected- and unaffected-side network components for S2 and S6 are presented below their corresponding statistical results. The relative muscle interaction strength (network edge widths), degree of involvement (node size) and submodular structure (node colour) are also illustrated. **(B)** The Canberra distance between the activation magnitudes at pre- and post-sessions separately for S3 and S4 significantly differentiated treatment responders and non-responders (Responders: 0.2±0.18, Non-Responders: 0.48±0.31 (t = 3.1 (*p* = 0.004)) (Median±SD)) and PT and VR treatment groups (PT: 0.19±0.18, VR: 0.33±0.26 (t = -2.2 (*p* = 0.044)) (Median±SD)) respectively. The affected- and unaffected-side network components for S3 and S4 is presented below their corresponding statistical results. The relative muscle interaction strength (network edge widths), degree of involvement (node size) and submodular structure (node colour) are also illustrated.

Examination of the extracted activation patterns relationship with treatment responsiveness among responders and non-responders and among PT and VR participants revealed further evidence for the crucial role of synergistic muscle interactions (Fig.2(B)). Specifically, we found that among non-responders S3 activation changed significantly after treatment than responders (t = 3.1 (*p* = 0.004)), while the VR group demonstrated a significant change in the activation of S4 compared to participants in the PT group (t = -2.2 (*p* = 0.044)). Interestingly, both S3 and S4 were functionally related to elbow flexion with shoulder girdle stabilisation (Fig.2(B) and Supplementary Materials Fig.2), suggesting that non-responders express maladaptive activation adjustments of specific flexor-driven networks, while VR treatment promotes changes in other flexor-driven networks.

### A redundancy-to-synergy transformation characterises effective rehabilitation

To characterise treatment responsiveness from the perspective of the NIF, we took the redundant and synergistic muscle networks of each participant on both affected- and unaffected-sides and statistically compared muscle network-level (i.e. average network interaction strengths) and interaction-level differences (i.e. individual muscle connection strengths (see ‘*Characterising rehabilitation effects on functional muscle interactions among responders and non-responders*’ section of Materials and Methods)) between sessions among conventionally-defined responders (i.e. >5 point change in FMA-UE scores) (Fig.3(A-B.1-2)) and non-responders (i.e. <5 point change in FMA-UE scores) (Fig.3(C-D.1-2)).

**Figure 3.**
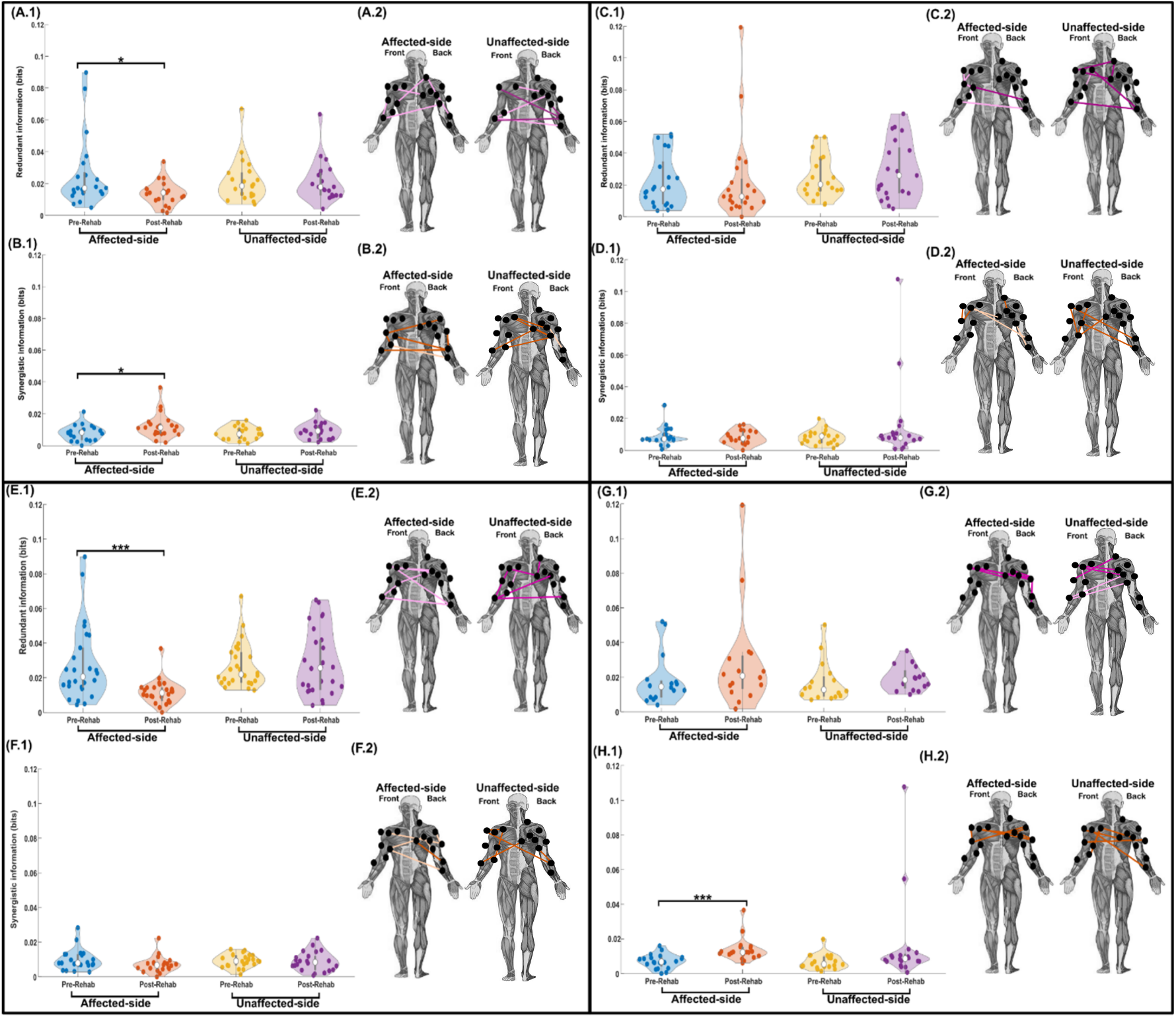
The average magnitudes of clinical assessment defined responders’ redundant and synergistic interactions (**A.1**-**B.1** respectively) and non-responders’ (**C.1**-**D.1** respectively also) for affected- and unaffected-sides at pre- and post-sessions are illustrated as violin plots. Significant decreases and increases (*p* < 0.05 (**\***)) from pre-post session were found on the affected-side for redundant and synergistic networks of responders respectively. To the right of the responder and non-responder plots (**A.2**-**B.2** and **C.2**-**D.2** respectively), muscle interactions identified to be significantly stronger at the pre-session (light coloured network edges) or at the post-session (dark coloured network edges) are illustrated for both affected- and unaffected-sides. For illustrative purposes, the 90th percentile of these significantly different muscle interactions are depicted only. Below, a corresponding output where participants were partitioned using *R*_Pre-Post_ (**E**-**F.1-2**) and *S*_Pre-Post_ (**G-H.1-2**). Pre-post differences here accentuated the differences found using the clinical assessment derived partition (**A-D.1-2**) with more significant decreases and increases in redundant and synergistic interaction strengths respectively (*p* < 0.001 (**\*\*\***)).

Treatment responders experienced a significant decrease in redundancy on the affected-side (t= 2.3 (*p* < 0.05) (Fig.3(A.1)) and significant increase in synergy on the affected-side also (t= -2.2 (*p* < 0.05) (Fig.3(B.1)). No noticeable changes were experienced for average redundancy or synergy on the unaffected-side across responders were observed (Fig.3(A-B.1)). Specific muscle interactions also reflected this trend for redundancy and synergy with rehabilitation among responders (see network edges in Fig.3(A-B.2)). Whereas for the corresponding muscle interactions on the unaffected-side (Fig.3(A.2)), a mixture of significantly greater and lower magnitudes from baseline were found. The most significant synergistic muscle interactions of responders’ affected-side were also predominantly greater at follow-up (Fig.3(B.2)), except for BRACH-SU which was lower at follow-up. This was also the case for the unaffected-side of responders (Fig.3(B.2)), where TEM-PT was the only muscle interaction of those depicted to be lower at follow-up.

Meanwhile non-responders demonstrated no significant changes in average redundancy or synergy from pre-to-post sessions on either affected- or unaffected-sides (Fig.3(C-D.1-2)). Several redundant and synergistic muscle interactions on the unaffected-side were found to increase from pre-to-post sessions (Fig.3(C-D.2)), suggesting rehabilitation may have influenced control of the unaffected-side among non-responders rather than the affected-side.

Altogether, these findings characterise post-stroke motor recovery as a redundancy-to-synergy information conversion, demonstrating quantitatively the enhanced functional re-differentiation of muscle interactions provided by effective rehabilitation.

### Data-driven patient clusters more precisely quantify treatment response patterns

Implementation of our clustering approach (see Fig. 6 and ‘*Clustering stroke survivors based on impairment severity and therapeutic (non-) responsiveness*’ section of the Materials and methods for details) to quantify treatment response patterns (i.e. pre-to-post session changes in modular activation patterns) identified two redundant and synergistic patient subgroups (*R*_Pre-Post_, *S*_Pre-Post_). Supplementary Materials Fig.3 illustrates these clusters which did not demonstrate significant differences (*p* > 0.05) for pre- and post-session FMA-UE scores and the change in FMA-UE scores across the intervention (ΔFMA-UE). However, direct comparison of *R*_Pre-Post_ and *S*_Pre-Post_ clusters to the FMA-based responsiveness classification in terms of network- and interaction-level differences across the clinical intervention (see ‘*Characterising rehabilitation effects on functional muscle interactions among (non-) responders*’ section of Materials and methods for further details) demonstrated a more precise quantification of treatment responsiveness.

More specifically, for *R*_Pre-Post_ (Cluster 1 (Fig.3(E.1-2)), Cluster 2 (Fig.3(G.1-2)), we found that cluster 1 participants significantly reduced redundancy on the affected-side from pre-to-post sessions (t= 3.5 (*p* < 0.001) (Fig.3(E.1)), while the unaffected-side experienced no significant changes. Several of the affected-side muscle interactions were found to decrease in strength at post-treatment, while the unaffected-side demonstrated mostly increased with rehabilitation (Fig.3(E.2)). Meanwhile, cluster 2 participants demonstrated a non-significant increase in the average redundancy of the affected-side and no perceived changes on the unaffected-side (Fig.3(G.1)). *S*_Pre-Post_ (Cluster 1 (Fig.3(F.1-2)), Cluster 2 (Fig.3(H.1-2)) demonstrated a significant increase in synergistic muscle interaction strengths among cluster 2 participants (t= -3.78 (*p* < 0.001) (Fig.3(H.1))), while no significant changes were found on the unaffected-side. The most significantly different synergistic muscle interactions on both affected- and unaffected-sides were all found to be greater at post-treatment (Fig.3(H.2)). On the other hand, for cluster 1 participants (Fig.3(F)), we found a non-significant decrease in synergy on the affected-side and no perceivable changes on the unaffected-side (Fig.3(F.1)). The most significant synergistic muscle interactions were mixed in both directions for cluster 1 participants affected-side and all greater at follow-up for the unaffected-side (Fig.3(F.2)).

Taken together, we found that the NIF-derived treatment response clusters not only closely aligned with but further accentuated the redundancy-to-synergy conversion underpinning conventional metrics (see ‘*Effective post-stroke rehabilitation as a redundancy-to-synergy information conversion*’ section of the Results), suggesting a more precise quantification of treatment responsiveness.

### Stroke survivors cluster into severely and non-severely impaired subgroups with distinct physiological responses to neurological insult

Separate cluster analysis of pre- and post session modular activation patterns across the post-stroke cohort (see Fig. 6 and ‘*Clustering stroke survivors based on impairment severity and therapeutic (non-) responsiveness*’ section of the Materials and methods for details) consistently revealed two redundant and synergistic patient subgroups at baseline (i.e. *R*_Pre_ and *S*_Pre_ (Fig.4(A-B))) and at followup (i.e. *R*_Post_ and *S*_Post_ (Supplementary materials Fig.4(A-B))). Table.1 below provides an overview of the patients assigned to each cluster identified, showing participants across treatment groups (VR vs PT) and both clinical assessment- and NIF-defined responders (R) and non-responders (NR) were present in each cluster.

**Table 1.** An overview of the patient clusters extracted using the NIF. The Mean ± Standard deviations for Age and Time from lesion onset are presented along with the number of patients from each treatment type (i.e. physical therapy (PT) or virtual reality (VR)), conventional treatment response classifications (i.e. responders (R) and non-responders (NR) defined by MCID thresholds on the FMA-UE), and the NIF-derived treatment response clusters (i.e. *R*_Pre-Post_, *S*_Pre-Post_).

| Patient details | $R_{Pre}$ | $R_{Post}$ | $S_{Pre}$ | $S_{Post}$ |
| --- | --- | --- | --- | --- |
| Age (Years) | C1=61.1±11.3 | C1=63.9±11.3 | C1=63.1±12.1 | C1=62.8±10.2 |
| | C2=63.4±11.5 | C2=60.9±13.1 | C2=62±12.5 | C2=61.9±14.5 |
| Time since lesion<br>(Months) | C1=9±19.8 | C1=12±18.9 | C1=7±10.6 | 4.5±3.9 |
| | C2=7.2±9.5 | C2=3.9±3.7 | C2=8.5±16 | 12.1±20 |
| Treatment type<br>(PT/VR) | C1=4/14 | C1=1/20 | C1=3/13 | C1=4/19 |
|  | C2=4/20 | C2=7/14 | C2=5/21 | C2=4/15 |
| Conventional classification<br>(R/NR) | C1=11/7 | C1=8/13 | C1=10/6 | C1=8/15 |
|  | C2=10/14 | C2=13/14 | C2=11/15 | C2=13/6 |
| NIF clusters<br>(R/NR) | C1=7/11 | C1=19/2 | C1=6/11 | C1=7/16 |
|  | C2=19/5 | C2=7/14 | C2=14/11 | C2=13/6 |

**Figure 4.**
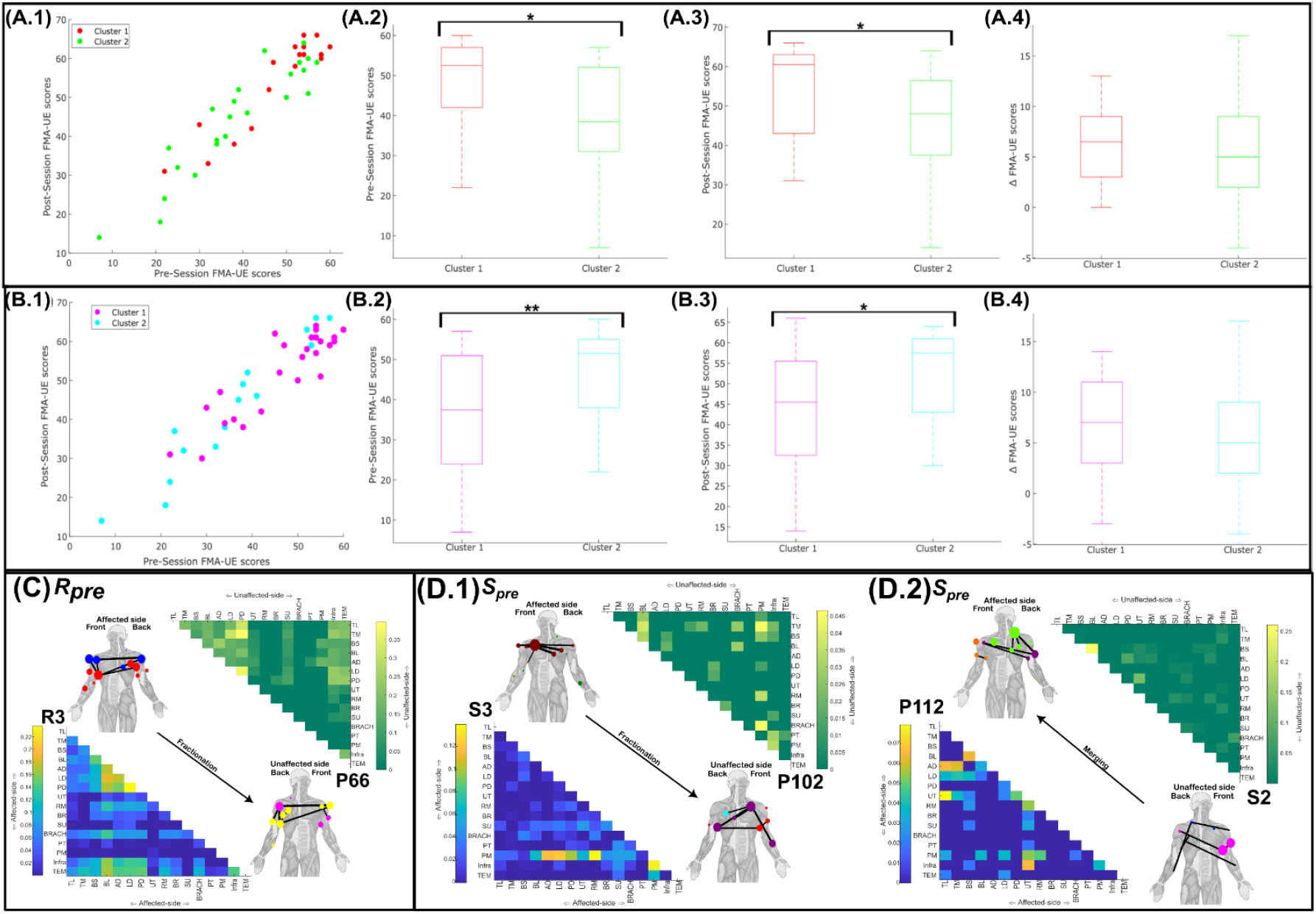
The identified patient clusters depicted with respect to pre- and post-session FMA-UE scores for *R*_Pre_ (**A.1**) and *S*_Pre_ (**B.1**). Boxplots illustrating the differences between the clusters identified in each partition for baseline FMA-UE scores (**A.2**-**B.2**), post-session FMA-UE scores (**A.3**-**B.3**) and the change in FMA-UE scores from baseline to follow-up (i.e. ΔFMA-UE) (**A.4**-**B.4**). **\*** indicates a significant difference of *p* < 0.05 and **\*\*** equates to *p* < 0.01. The network components identified as significantly contributing to (**C**) *R*_Pre_ and (**D.1-2**) *S*_Pre_ through fractionation (affected-s de (lower triangular matrix)) and merging (unaffected-side (upper triangular matrix)). For interpretation, a corresponding network from a representative participant accompanies each significant network component (i.e. P66, P102 and P112). The submodular structure (node colour) and most proportionally significant (i.e. >95th percentile) muscle interactions (network edges) are also illustrated for each network on human body models. (**C**) Fractionation of R3 explained participants’ affiliation with cluster 1 of *R*_Pre_ (*β* = -5.84±2.23 (*p* < 0.01)), classifying 78.6% of participants correctly. (**D.1**) Fractionation of S3 (*β* = -34.82±10.9 (*p* = 0.001)) and (**D.2**) merging of S2 (*β* = -16.4±8.6 (*p* = 0.056)) explained participants affiliation with *S*_Pre_ cluster 1, together classifying 71.4% of participants correctly. The patient clusters for *R*_Post_ and *S*_Post_ along with the main network components and associated physiological response patterns underpinning them are presented in the Supplementary materials Fig.4.

A significant difference (**\***) between clusters was found for *R*_Pre_ for both pre-session FMA-UE (t= 2.27 (*p* = 0.029) (Fig.4(A.2))) and post-session FMA-UE (t= 2.37 (*p* = 0.023) (Fig.4(A.3))), but not for the change in FMA-UE score from pre-to-post sessions (ΔFMA-UE) (Fig.4(A.4)). The *S*_Pre_ partition also significantly differentiated participants based on motor impairment at baseline (t= -2.9, *p* < 0.01(**) (Fig.4(B.2))) and follow-up sessions (t= -2.5, *p* = 0.017(**\***) (Fig.4(B.3)), but similarly to *R*_Pre_, could not distinguish participants based on ΔFMA-UE (Fig.4(B.4)). For both *S*_Post_ and *R*_Post_, presented in Supplementary materials Fig.4(A-B) respectively, the extracted partitions could not significantly differentiate participants in terms of pre- or post-session FMA-UE scores or ΔFMA-UE.

To shed mechanistic light on the underlying contributions to the patient clustering’s observed in Fig.4(A-B) and Supplementary materials Fig.4(A-B) here, we sought to determine if they captured distinct pattern of merging, fractionation and preservation among the post-stroke cohort (see ‘*Quantifying the contributions of preservation, merging and fractionation to impairment-based patient clusters*’ section of Materials and methods for details on their quantification). The network components and their corresponding physiological responses found to be significantly different between *R*_Pre_ and *S*_Pre_ patient clusters are illustrated in Fig.4(C-D) while the corresponding outputs for *R*_Post_ and *S*_Post_ are presented in Supplementary materials Fig.4(C-D). For interpretation, alongside each significantly contributing module, we have illustrated a corresponding network from a representative participant involved in each calculation.

To summarise, we found that for *R*_Pre_ (Fig.4(C)), the prevalence of fractionation in R3 (comprised of interactions among the shoulder prime-movers (i.e. PD, AD and LD)) was associated with the less impaired cluster 1 participants (*β* = -5.84±2.23 (*p* < 0.01), 78.6% classification accuracy). This evidence suggests that the fractionation of specific redundant modules is an adaptive response to neurological insult among non-severely impaired stroke survivors.

Fractionation of S3 along with merging of S1 at baseline contributed to the patient clustering’s observed in the more severely impaired subgroup of *S*_Pre_ (Fig.4(D.1-2)), with both associated with cluster 1 patients (*β* = -34.82±10.9 (*p* = 0.001), *β* = -16.4±8.6 (*p* = 0.056) respectively (Classification accuracy= 83.3%)). The fractionation of muscle groups involving the shoulder girdle (i.e. PM (Fig.4(D.1))) was again an important contributor to the observed patient groupings, however in contrast to the redundant modules (Fig.4(C)), here fractionation of synergistic modules was associated with the severely impaired patient cluster. The first instance of merging patterns contributing to patient subgroupings here was found for *S*_Pre_, demonstrating how the maladaptive merging of elbow flexor related interactions (i.e. BL-BS (S1) (Fig.4(D.2)) underpins the more severely impaired stroke survivors.

Fractionation of R4 (centralised around redundant interactions between UT and both scapula stabilisers and wrist extensor muscles) at follow-up was also associated with the cluster 1 participants of *R*_Post_ (*β* = -2.32±0.99 (*p* = 0.019) (Supplementary materials Fig.4(C.1))). The underlying contributions to *R*_Post_ clusters were more complex than *R*_Pre_ however, including an additional contribution by the preservation patterns in R1 (an elbow extensor related functional module) (*β* = -15.2±6.5 (*p* = 0.018)) (Supplementary materials Fig.4(C.2))). Together, these contributions illustrate that cluster 1 participants of *R*_Post_ had significantly greater fractionation of R4 and preservation of R1 compared to cluster 2 participants at follow-up.

Finally, patterns of fractionation (S1 (Supplementary materials Fig.4(D.1)) and of merging (S2 (Supplementary materials Fig.2(D.2))) were found to significantly contribute to the *S*_Post_ clusterings, both associated with cluster 2 affiliation (*β* = 26.7±12.1 (*p* = 0.027) and *β* =27.3±11.8 (*p* = 0.021) respectively (Classification accuracy: 76.2%)). S1 here, like S3 in *S*_Pre_ (Fig.4(D.1)), comprised of PM playing a central role in interaction with all other muscles (Supplementary materials Fig.4(D.1)). There wasn’t however a clear correspondence between the contributing merging patterns for *S*_Pre_ and *S*_Post_ (Fig.4(D.2) and Supplementary materials Fig.4(D.2) respectively). Instead, for *S*_Post_ the merging pattern significantly differentiating participants at post-treatment predominantly involved the SU.

## Discussion

This study leveraged a novel muscle network analysis framework to provide a data-driven account of post-stroke motor impairment and therapeutic responsiveness and, in doing so, address current challenges to the standardised assessment of stroke survivors. Multiple flexor- and extensor-driven modules were uncovered from a post-stroke cohort undergoing intensive upper-limb motor training, collectively representing the diverse biomechanical contributions of functional muscle networks to task performance. Their activation patterns were closely associated with gross motor function impairment and both treatment type and responsiveness, promoting their use as clinical biomarkers. Deployment of a novel clustering algorithm revealed consistent dichotomous patient subgroupings reflecting clinical assessment-based classifications of motor impairment (i.e. severely vs. non-severely impaired patients) and treatment responsiveness (i.e. responders and non-responders) underpinned by distinct physiological markers of impairment (i.e. muscle network merging, fractionation, and preservation). Crucially, we provided a novel characterisation of effective post-stroke rehabilitation as a redundancy-to-synergy information conversion that highlights functional re-differentiation of muscle interactions with effective treatment, a response pattern closely reflected in conventional clinical assessment-based classifications and further accentuated in the derived patient clusters. Our work here provides novel neurobiological insights into and biomarkers of movement pathology whilst offering a principled approach for precision motor assessments in stroke survivors.

### Altered composition and activation of functional muscle networks post-stroke

The functional characterisation of muscle interactions as redundant (i.e. functionally-similar) and - synergistic (i.e. functionally-complementary) is a unique capability of the NIF (O’Reilly & Delis, 2024). By extracting these distinct types of functional interaction, we uncovered multiple task-relevant extensor- and flexor-driven synergies that, remarkably, were found predominantly among redundant and synergistic muscle networks respectively (see Supplementary Materials Fig.1-2). These findings align with recent work demonstrating multiple modules underpin the post-stroke flexor synergy (Kim et al., 2024a), going further in showing that multiple modules also underpin the post-stroke extensor synergy and that both synergies are comprised predominantly of distinct types of functional interaction. Continuing, the activation of several flexor-driven synergistic muscle networks closely reflected motor impairment consistently across the intervention and both treatment-type and -responsiveness (Fig.2), exemplifying the central role of the flexor synergy in manifestations of functional impairment post-stroke (Alt Murphy et al., 2022; Ellis et al., 2017). We present the NIF here as a principled methodology for the separate quantification of abnormal flexor- and extensor-synergy expression along with their distinct contributions to motor function and treatment responsiveness.

Impaired control of not only prime-mover muscle groups but also secondary, stabilising musculature are prevalent post-stroke (Paci et al., 2007; Ryerson et al., 2008; Silva et al., 2018). In contrast to traditional muscle synergy analysis (Bruton & O’Dwyer, 2018; Cheung & Seki, 2021; Turpin et al., 2021), the NIF does not rely on optimising data variance explained and therefore the functional contributions of more subtle interactions often marginalised in traditional muscle synergy approaches were equivalently emphasised alongside higher amplitude muscle covariations (see Supplementary Materials Fig.1-2) (O’Reilly & Delis, 2024; Ó’Reilly & Delis, 2022). This additional capability revealed widespread redundant and synergistic interactions among secondary stabilising musculature (e.g. teres major, infraspinatus and rhomboids) and with prime-mover muscles (e.g. triceps and biceps brachii, deltoids) that frequently demonstrated high network prominence and interaction strengths (see Tables in Supplementary Materials Fig.1-2). Hence, the NIF offers novel clinical utility to the assessment of these subtle muscle interactions underlying motor functions commonly impaired post-stroke such as anticipatory postural adjustments and segmental/joint stabilisation (Paci et al., 2007; Silva et al., 2018).

The fractionation of motor patterns into separate modules in response to neurological insult emerged as a central difference between the patient clusters identified here (see Fig.4 and Supplementary materials Fig.4). We found new evidence for the adaptive role of fractionation among redundant muscle interactions but maladaptive role of fractionation among synergistic muscle networks (Fig.4). This provides evidence towards crucially unresolved questions on the role of muscle synergy fractionation from the original work presenting these physiological markers (Cheung et al., 2012), demonstrating that it indeed depends on the type of functional muscle interaction involved. Further, the NIF demonstrated here an enhanced capability over traditional approaches to identify these crucial patterns, as earlier work on related versions of this dataset could not identify any differentiable fractionation events across the cohort (Pregnolato et al., 2025). Although fractionation was found to also differentiate patient clusters at follow-up (Supplementary materials Fig.4(C-D)), the exact role fractionation played in treatment responsiveness was unclear here as no significant differences between post-session patient clusters in terms of FMA-UE were observed (Supplementary materials Fig.4(A&B.3). Finally, the observations of both the merging and preservation of motor patterns contributing to gross motor function impairment and perhaps treatment responsiveness aligns with the existing literature (Cheung et al., 2012; Clark et al., 2010; García-Cossio et al., 2014; Hashiguchi et al., 2016), and offers the potential for additional insight into their frequently contrasting effects on motor function depending on the individual’s condition—whether it’s impairment severity, developmental stage, or training experience (Cheung et al., 2020; García-Cossio et al., 2014).

### Post-stroke motor recovery as a redundancy-to-synergy information conversion

In this study, we provided a novel characterisation of treatment-induced motor recovery among stroke survivors as a redundancy-to-synergy conversion of task information (Fig.3). Specifically, a reduction in redundancy and increase in synergy on the affected-side of responders was found while no such changes were observed among non-responders or on the unaffected-side. This pattern was not only observed in the functional muscle networks of patients grouped in the conventional way using the MCID of the FMA-UE measure, but also in the NIF-derived patient clusters (i.e. *R*_Pre-Post_, *S*_Pre-Post_) where it was in fact further accentuated. Previous research has shown that functional redundancy increases post-stroke (Cheung et al., 2012; Clark et al., 2010), reflecting the characteristic loss of functional specificity (i.e. functional de-differentiation) of muscle interactions post-stroke. Enhanced synergy with treatment here thus reflects the functional re-differentiation of predominantly flexor-driven muscle networks towards different, complementary task-objectives across the seven upper-limb motor tasks performed (Kim et al., 2024b), leading to improved motor function among responders. Although the exact neural mechanisms underpinning redundant and synergistic muscle interactions remain a future research matter, abnormal muscle interactions post-stroke are known to originate from damage to the corticospinal tract (CST), particularly effecting flexor muscle groups and which are compensated for by reticulospinal tract (RST) upregulation (García-Cossio et al., 2014; Kim et al., 2024a; McPherson et al., 2018). Both neural pathways are known to exhibit specific flexor-extensor biases (Baker, 2011), but both indeed provide parallel input to motoneurons to coordinate groups of muscles towards complementary functional objectives (e.g. fine vs. gross motor functions) (Glover & Baker, 2022; Sangari & Perez, 2020). Hence, treatment-induced enhancement of functional complementarity among predominantly flexor-driven muscle networks here could reflect CST functional recovery and therefore, a reduced reliance on alternative neural pathways to execute the same function (Kim et al., 2024a, 2024b; McPherson et al., 2018). These observations promote the clinical use case of NIF in the precise assessment and monitoring of neural impairment over traditional approaches (O’Reilly & Delis, 2024; O’Reilly & Delis, 2025; O’Reilly & Delis, 2024).

### Dichotomous clustering of motor impairment and treatment responsiveness in the post-stroke population

Through the novel clustering of stroke survivors presented here, we consistently identified dichotomous subgroupings that reflected distinct patterns of motor impairment and treatment responsiveness (Fig.3-4 and Supplementary materials Fig.3-4). These clustering’s are formed through the greater affinity of some stroke survivors compared to others in terms of their modular activations and changes thereof with treatment. Based on our own and others’ previous work (García-Cossio et al., 2014; O’Reilly & Delis, 2024; O’Reilly & Delis, 2025), this functional grouping results from the preservation of unimpaired modules among certain cohorts and the generation of new motor patterns in others, a distinction seemingly dependent on residual CST function (García-Cossio et al., 2014). These findings link well also with the increasing awareness in the rehabilitation literature for the distinct aetiology of severe and non-severe impairments in stroke survivors that is not merely an artefact of measurement or unfounded simplification (Bonkhoff et al., 2022; Byblow et al., 2015; García-Cossio et al., 2014; Kim et al., 2024a; O’Reilly & Delis, 2024). This awareness comes from the distinct rates of proportional recovery observed among stroke survivors with and without an intact CST (Bonkhoff et al., 2022; Byblow et al., 2015). These observations do not rule out the possibility of more fine-grained patient subgroups, as preliminary applications without sparsification and related work employing the clustering algorithm employed here revealed 5 further subclusters, aligning with previous research (O’Reilly & Delis, 2025; Scano et al., 2017; van der Vliet et al., 2020). Indeed, in future work we aim to apply manifold learning approaches to the co-membership matrix derived from this clustering algorithm, providing a continuous representation of the population structure. Nevertheless, the principled quantification of more generally applicable yet crucially discernible patient clusters offers new opportunities in the rehabilitation of severely impaired stroke survivors, a cohort that experiences significantly greater challenges during rehabilitation (Angerhöfer et al., 2021; McGlinchey et al., 2020).

## Limitations

Although the FMA-UE is a gold standard measure of post-stroke treatment outcomes (Meyer et al., 1975; Page et al., 2012), it does not capture the impact of rehabilitation on patients’ ability to perform activities-of-daily-living or to participate in daily life. Hence, interpretations of the identified biomarkers are currently limited to gross motor function impairment and recovery. Future research should employ this framework to quantify biomarkers that correspond to other important aspects of patients’ recovery (e.g. functional independence, subjective experiences), thus offering a more complete evidence base for its clinical utility.

## Methods and Materials

### Experimental setup and Data collection

We conducted a secondary analysis on previously published data (Maistrello et al., 2021; Pregnolato et al., 2025) (ethical approval provided by the local Ethical Committee of the IRCCS San Camillo Hospital s.r.l, Venice, Italy), including 42 stroke survivors (Age (Years): 62.4 ±12.2, Gender (Male/Female): 28/14, Diagnosis (ischaemic/haemorrhagic): 36/6, Time from lesion onset (Months): 7.5±14.1). All participants had previously been hospitalised with their first stroke. Exclusion criteria for this study included: 1) Mini Mental State Examination (MMSE) score lower than 20 points indicating moderate cognitive impairment; 2) severe verbal comprehension deficits, defined >13 errors on the Token Test; 3) evidence of apraxia and visuospatial neglect, evaluated by neurological examination; and 4) history of behavioural disorders (e.g., depression, aggressiveness, apathy) or neurological/vascular co-morbidity. This pre-post longitudinal study involved 4 weeks of intensive treatment consisting of 20 sessions of upper-limb exercises (one hour/session, 5 sessions/week). Participants that performed less than 80% of the planned sessions (< 16/20 sessions) were considered as study drop-outs and not included in further analyses. Two treatment groups were defined for this study which participants were randomly assigned into either conventional physical therapy (PT, No. participants enrolled= 8) or virtual reality (VR, No. participants enrolled=34) intervention (VRRS®, Khymeia Group Ltd., Noventa Padovana, Italy) groups (see the parent papers of this experimental protocol for further details (Maistrello et al., 2021; Pregnolato et al., 2024)) (Fig.1(B)). The Fugl-Meyer Assessment Upper Extremity section (FMA-UE), a gold standard clinical measure for assessing gross motor functions was used to quantify motor impairment in the affected upper-limb (Meyer et al., 1975), both prior to and immediately following the 4-week intervention. A minimal clinically important difference (MCID) of 5 points change at FMA-UE scores after treatment was used to classify responders and non-responders in both groups (Page et al., 2012). Moreover, before and after treatment an instrumental assessment was conducted recordings sEMG (EMG-USB2+, OT Bioelettronica, sampling rate: 2000Hz) from 16 muscles (triceps -medialis (TM) and lateralis (TL); biceps short head (BS) and biceps long head (BL); anterior deltoid (AD); lateral deltoid (LD); posterior deltoid (PD); upper trapezius (UT); rhomboid major (RM); brachioradialis (BR); supinator (SU); brachialis (BRACH); pronator teres (PT); pectoralis major (clavicular head) (PM); infraspinatus (Infra); teres major (TEM) (Fig.1(C))) of both affected and unaffected sides, while participants performed 10 repetitions of seven standardised motor tasks (Table.2). To prepare the sEMG signals for subsequent analyses, they were full-wave rectified and low-pass filtered at 20Hz using a 4th-order, bi-directional Butterworth filter with zero-phase distortion.

**Table 2.** The seven tasks performed by each participant before and after the clinical intervention including descriptions of starting position and movement involved.

| Task | Starting Position | Movement |
| --- | --- | --- |
| Simple Reaching | Hand fully pronated with lightly closed fist on the same side lap at a position close to the knee. | Move hand vertically along a straight path to eye level, fully extending and pronating elbow and wrist respectively. |
| Abduction | Hand fully pronated with lightly closed fist on the same side lap at a position close to the knee. | Move hand laterally to the shoulder level by abducting and flexing shoulder while keeping elbow and wrist fully extended and pronated throughout respectively. |
| Reaching with single spatial constraint | Hand fully pronated and with lightly closed fist at the midpoint position between knee and hip on the same side lap. | Move hand to shoulder level in front with elbow extension maximal by trial end. Most of the elbow extension should occur after movement midpoint. |
| Reaching with two spatial constraints | Hand fully pronated with lightly closed fist on same side lap and close to the knee. | This task was identical to task 1 but with more movement constraints. |
| Pronation | Shoulder should be adducted, elbow neutral at 90°, wrist fully supinated and shoulder laterally rotated so the forearm is approximately 45° relative to the thigh. | Follow a circular movement trajectory where the wrist pronates and is placed on the same side lap. |
| "Cactus" | Full elbow extension and shoulder adduction and extension with lateral alignment. Wrist can stay pronated. | Full shoulder abduction, elbow extension and wrist pronation throughout. Return to starting position. |
| Pronation with a path | Full elbow extension and wrist supination. Shoulder abduction approximately 45° relative to the trunk. | Full shoulder adduction and elbow extension throughout while wrist smoothly pronates to rest on the same side lap. |

### Extraction of Redundant and Synergistic Muscle Networks

To quantify the task-relevant information shared by interacting muscles and then characterise them as having either functionally-similar (i.e. redundant) or -complementary (i.e. synergistic) roles, we employed the NIF to map networks of muscle interactions to task performance and extract the underlying low-dimensional components (O’Reilly & Delis, 2024). More specifically, leveraging a higher-order information-theoretic measure known as co-information (*II*) (Ince et al., 2017; McGill, 1954) (Fig.1(A)), we firstly quantified the task-relevant information (pink-orange intersection) shared between all possible muscle pairings (e.g. [*m*_*x*_, *m*_*y*_]) with respect to a given task parameter (*τ*), formally disentangling it from the task-irrelevant space (yellow intersection) (O’Reilly & Delis, 2024). *II* quantifies the balance between redundancy (i.e. the task-specific variations that are equivalently shared by *m*_*x*_ and *m*_*y*_ and so provide less information when combined) and synergy (i.e. the task-specific variations of *m*_*x*_ and *m*_*y*_ that when observed together result in additional predictive information about *τ*) by quantifying the difference between the task information shared (i.e. the mutual information (Ince et al., 2017)) provided by *m*_*x*_ and *m*_*y*_ separately, *I*(*m*_*x*_; *τ*) + *I*(*m*_*y*_; *τ*), and shared together, *I*(*m*_*x*_, *m*_*y*_; *τ*) (Fig. 5). Consequently, the functional role of each muscle interaction can be principally quantified as net redundant (negative *II*; pink shading of pink-orange intersection) or synergistic (positive *II*; orange shading of pink-orange intersection). These distinct quantities generalise the conceptual underpinning of muscle synergies (Bruton & O’Dwyer, 2018), formally distinguishing between two alternative ways in which muscles may ‘*work together*’ towards task performance.

**Figure 5.**
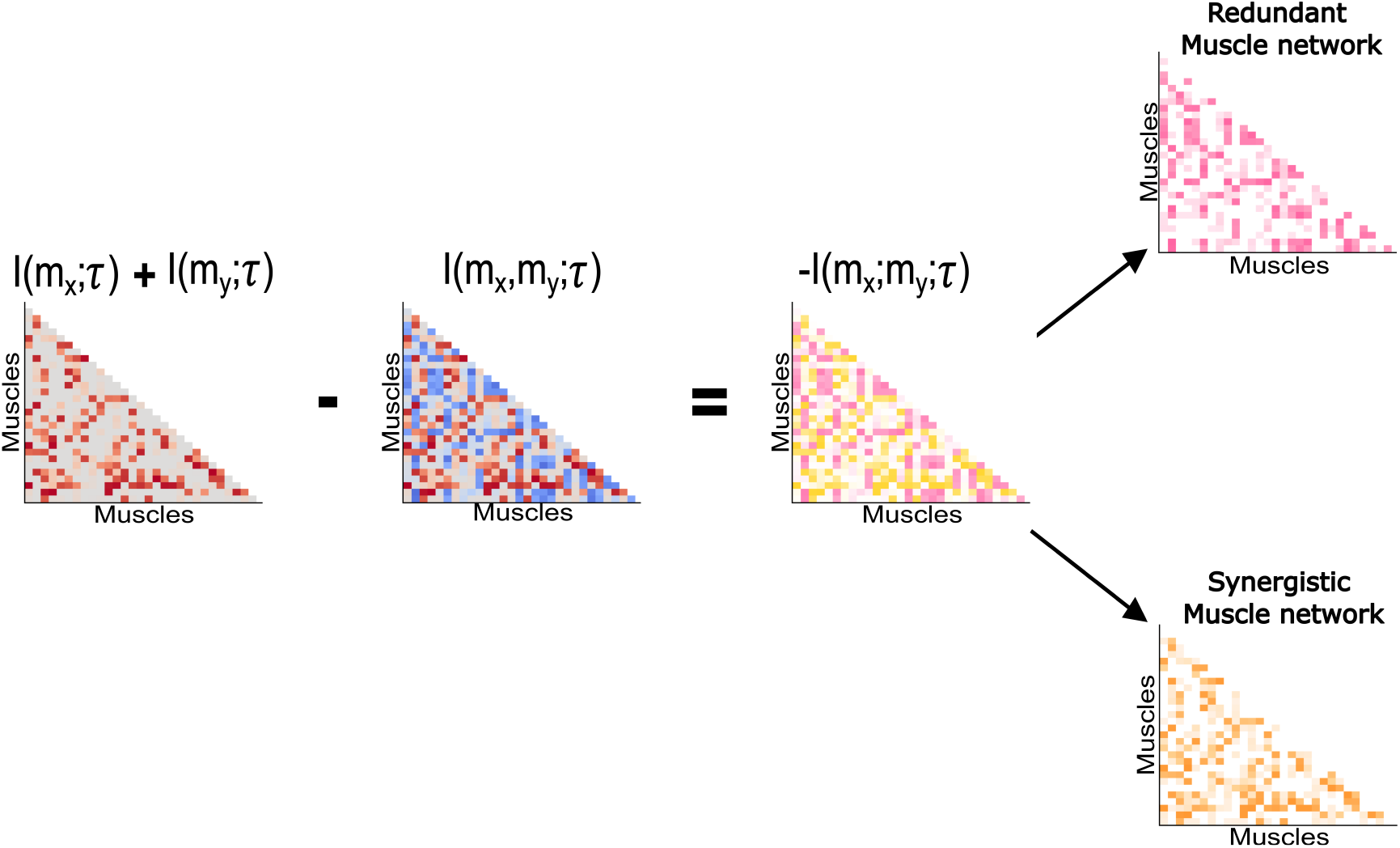
*II* (−*I*(*m*_*x*_; *m*_*y*_; *τ*)) determines the difference between the sum of mutual information with in *m*_*x*_ and *m*_*y*_ when observed separately (*I*(*m*_*x*_; *τ*) + *I*(*m*_*y*_; *τ*)) and the mutual information with when observed together (*I*(*m*_*x*_, *m*_*y*_; *τ*)). The corresponding adjacency matrices (*A*) show how this calculation is carried out collectively for all unique [*m*_*x*_, *m*_*y*_] pairings. Net redundant (pink) and synergistic (orange) muscle couplings are then separated into sparse, non-negative networks.

For this study’s analysis, we generated redundant and synergistic networks from the affected and unaffected side muscles separately with respect to a discrete task parameter describing the seven motor tasks each participant performed. We carried out this application in the spatial domain (i.e. interactions between muscles across time (Ó’Reilly & Delis, 2022)) by concatenating the 10 repetitions of each task executed on a particular side (i.e. variables of length no. of timepoints x 10 trials) and quantifying *II* with respect to this discrete task parameter codified to describe the motor task performed at each timepoint for each trial included. This computation was performed on all unique *m*_*x*_ and *m*_*y*_ pairings, generating symmetric matrices (*A*) (i.e. *A* = *A*^*T*^) composed separately of non-negative redundant and synergistic values (Fig. 5). Having quantified these distinct types of task-relevant muscle networks for each participant on both affected and unaffected sides, the following steps in the NIF pipeline were then performed:

- **Step 1: Network sparsification** We empirically sparsified each *A* separately by identifying robust muscle network structures not found in equivalent random networks using a modified percolation analysis (Gallos et al., 2012), setting non-significant muscle interactions to zero.
- **Step 2: Model-rank specification** We then determined the functional organisation of each *A* using a link-based community detection method (implemented with single linkage) (Ahn et al., 2010) (Fig. 2(C.1)), essentially unravelling the nested structure of redundant or synergistic interactions into separate co-membership matrices for each module identified (i.e. entries indicate where a pair of nodes are functionally affiliated or not) (O’Reilly et al., 2025; O’Reilly & Delis, 2024). A consensus partition across participants was then subsequently determined by aggregating across all co-membership matrices and then applying the Louvain algorithm to the resultant matrix (Blondel et al., 2008; Rubinov & Sporns, 2010).
- **Step 3: Dimensionality reduction** The number of modules identified in this final consensus partition of step 2 was used as the input parameter for dimensionality reduction, namely projective non-negative matrix factorisation (PNMF) (Fig. 1(D)) (Yang & Oja, 2010). The choice of PNMF here which has been employed in earlier work on the NIF (O’Reilly & Delis, 2022), in contrast to the space-time tensor decomposition employed in the parent study (O’Reilly & Delis, 2024), was chosen simply to maintain brevity by focussing the analysis on the spatial domain. Continuing, the input matrix for PNMF consisted of the sparsified *A* on both affected and unaffected sides from all participants at both pre- and post-sessions concatenated together in their vectorised form. More specifically, the input matrix composed of redundant or synergistic values was configured such that the set of unique muscle pairings (1… *K*) on affected and unaffected sides (*x*_aff_ and *x*_unaff_ respectively) corresponding to an individual participant (*p*) from the group of participants (1… *N*) are concatenated row-wise and then the set of pre-session (pre) and post-session (post) networks are concatenated column-wise (Equation 1.1). Factorisation of this matrix results in the extraction of, for the *j*th module, a representative set of co-occurring muscle interaction weightings (*v*) across affected and unaffected sides and their underlying session- and participant-specific activation coefficients (*a*).

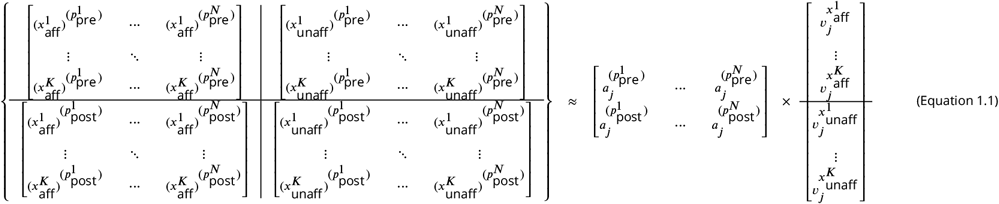

The extracted components can be reshaped into fully connected adjacency matrices (i.e. muscle networks) where the magnitude of the edges represents the interaction strength between muscle pairs. For the purpose of illustrations, the most prominent muscle interactions (i.e. proportional thresholding (*p* < 0.05)) within each extracted module are presented as a network over a human body model (Rubinov & Sporns, 2010). To visualise the extracted networks’ submodular structure, we applied the Louvain algorithm and depicted the partitions as distinct node colours on a human body model (Blondel et al., 2008; Rubinov & Sporns, 2010). Moreover, to illustrate the principal muscles involved in each component, the interaction strengths and relative importance of individual muscles were used to indicate edge-widths and node sizes respectively. Relative importance of muscles was quantified via the eigenvector centrality (i.e. the eigenvector associated with the largest eigenvalue in the network) (Newman, 2008), where muscles were considered prominent when they were well-connected with other well-connected muscles. Matlab code for this analytical pipeline has been published here: https://github.com/DelisLab/EMG2Task. To facilitate comparisons with existing approaches, we performed a conventional muscle synergy analysis on the post-stroke cohort (see Supplementary Materials Fig. 5 and associated text). Further comparisons with conventional approaches can be found in our previous work developing this framework (O’Reilly & Delis, 2022; O’Reilly & Delis, 2024).

### Clustering stroke survivors based on gross motor impairment severity and therapeutic responsiveness

To understand how the stroke survivors compared in terms of gross motor impairment severity and treatment-responsiveness using NIF, we conducted cluster analyses using the extracted activation coefficients (*a*). We focussed on *a* here as the extraction of population-level functional modules enabled the buffering of individual differences into the space of modular activations, making them an ideal target for identifying population structure. Towards finding an effective approach to clustering participants in this data based on differences in impairment severity and therapeutic (non-)responsiveness, we found that conventional clustering algorithms (e.g. agglomerative, k-means etc.) could not provide substantive outputs (see Supplementary Materials Fig. 5 and associated text for a direct comparison with conventional approaches), perhaps resulting from the complex interdependencies between the modular activations. Therefore, we sought to improve the linear separability of the participants by mapping *a* into a higher-dimensional feature space, and then applying network community detection protocols already incorporated into the NIF here to discern the underlying population structure (Fig. 6) (O’Reilly & Delis, 2025). This mapping was performed separately at pre- and post-sessions for motor impairment and from baseline to follow-up for treatment responsiveness (Fig. 6(A.1-2)). To carry out this feature mapping, we conducted a grid-search over different kernel functions (*K*) (i.e. linear, Gaussian, and polynomial functions) and identified parsimonious clusterings.

**Figure 6.**
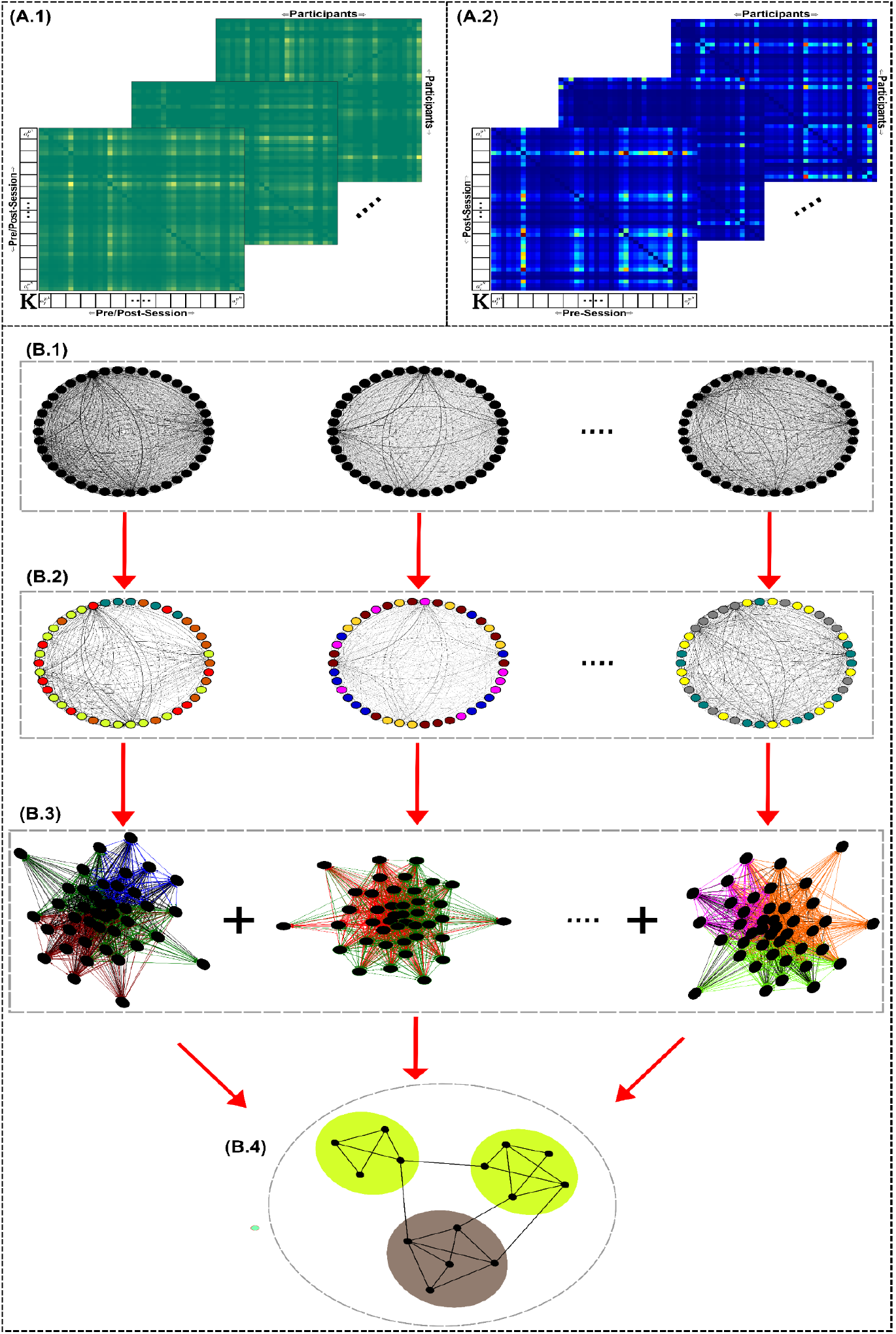
An overview of the divisive clustering algorithm employed (O’Reilly & Delis, 2025). Kernel matrices were computed from all possible pairwise combinations of functionally-redundant or -synergistic modular activations, generating a multiplex network (*C*) where each layer represents the similarity between all participant pairings for a given pair of module activations. (**A.1**) To cluster participants based on motor impairment, this meant computing C using either the pre- or post-session activations separately, (**A.2**) while for treatment responsiveness this computation was carried out between pre- and post-session modular activations. The dense layers of *C* (**B.1**) were empirically sparsified and their modular structure was quantified using a community detection protocol (the node colours represent different network communities) (**B.2**). (**B.3**) Co-membership matrices were generated from each layer-specific computation (i.e. entries indicate whether a participant pairing belong to the same cluster or not) which were then subsequently aggregated into a single representative matrix. (**B.4**) Finally, network community detection was re-applied to this representative matrix to quantify patient clusters.

We found that for distinguishing participants’ motor impairment level using redundant modular activations (*R*_Pre_, *R*_Post_), the linear kernel (i.e. 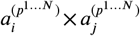) was most effective, while for synergistic activations (*S*_Pre_, *S*_Post_), the Gaussian kernel (i.e. exp 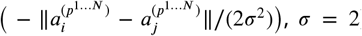) was best suited for distinguishing participants’ motor impairment. Finally, for treatment responsiveness (*R*_Pre-Post_, *S*_Pre-Post_), we found that the linear kernel provided a meaningful participant clustering. In the following, we briefly outline the steps involved in this clustering algorithm (Fig. 6):

- **Step 1: Multiplex feature mapping** Local kernel matrices were generated from each pairwise combination of extracted module activation coefficients across all participants (i.e. 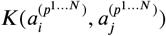) (see Fig. 6(A.1) for clustering based on motor impairment and Fig. 6(A.2) for clustering based on treatment responsiveness). This results in a multiplex network (*C*) comprised of layers (*c*_*i*_) for each module combination that described the similarity of each pair of participants’ modular activation (Fig. 6(A.1)). For impairment-based clustering (Fig. 6(A.1)), the diagonal of *c*_*i*_ (representing the self-similarity of each participant’s modular activation) was set to zero (Murphy et al., 2018), while for treatment responsiveness clustering the diagonal was maintained as it provided important self-referential information regarding the changes in participants’ modular activation patterns across the intervention (Fig. 6(A.2)).
- **Step 2: Sparsification and patient clustering** The densely connected *c*_*i*_ (Fig. 6(B.1)) were then empirically sparsified using a modified percolation analysis before their modular structures were separately quantified using the Louvain algorithm (node colours in Fig. 6(B.2) represent different network communities) (Blondel et al., 2008; Gallos et al., 2012; Rubinov & Sporns, 2010). To offset any stochasticity in the solutions found in the final application of the Louvain algorithm, we re-applied it *n* = 5000 times and determined a consensus partition using an established method (Lancichinetti & Fortunato, 2012). Following this, co-membership matrices were yielded from all *c*_*i*_ partitions, where entries indicate whether a participant pairing belong to the same cluster or not (Fig. 6B.3)). These co-membership matrices were then subsequently aggregated into a single representative matrix (Fig. 6(B.3)).
- **Step 3: Patient cluster identification**. Finally, the Louvain algorithm is re-applied to this representative matrix to uncover patient clusters (Fig. 6(B.4)).

Matlab code for this clustering algorithm has been published here: https://github.com/DelisLab/NICF.

### Quantifying the contributions of preservation, merging and fractionation to impairment-based patient clusters

To investigate whether the distinct patterns of preservation, merging and fractionation underlie the identified patient clusters in terms of motor impairment at pre- and post-sessions separately (i.e. [*R*_Pre_, *R*_Post_], [*S*_Pre_, *S*_Post_]), we implemented the following analysis.

To identify patterns of merging and fractionation among the functionally-similar and -complementary muscle networks of this post-stroke cohort, we aimed to identify linear combinations of representative affected- and unaffected-side muscle networks extracted across participants that could reconstruct individual participants’ muscle networks. More specifically, we concatenated all participants’ affected and unaffected-side redundant/synergistic muscle networks in a similar way as described in Equation 1.1 (see *‘Extraction of Redundant and Synergistic Muscle Networks’* section of Materials and Methods) but separately for pre- and post-session data. We then determined the optimal number of components to extract for each session across participants using PNMF (see steps 2–3 of ‘*Extraction of Redundant and Synergistic Muscle Networks*’ from the Materials and Methods section).

To quantify merging events for each participant at baseline and follow-up sessions, we determined the weighted contributions of the extracted unaffected-side network components (v^*u*^) to the reconstruction of the *p*th participant’s affected-side network 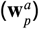 using a non-negative least squares algorithm (‘*lsqnonneg*’ Matlab function) (Equation 2.1) (Cheung et al., 2012). Here, the non-negative, participant-specific merging coefficient (*m*) describes the contribution of the *k*th unaffected-side component from the set of extracted session-specific network components (*N*) to the reconstruction of each individual participant’s network:

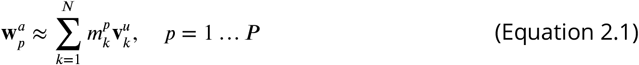

In a similar vein, fractionation was quantified by essentially reversing this computation to determine participant-specific fractionation coefficients (*f*) for the *k*th affected-side network component extracted (v^*a*^) from individual participants’ unaffected-side muscle network 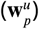 (Equation 2.2):

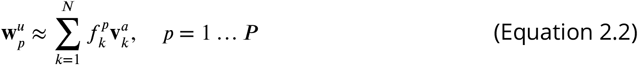

To determine patterns of preservation for the *p*th participant at baseline and follow-up sessions, we quantified the inverse of the Euclidean distance as a measure of similarity (*s*) between the *k*th v^*u*^ and 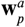 (Equation 2.3). Here, 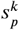 ranges from 0 to 1 with values close to 1 indicating perfect similarity between the networks:

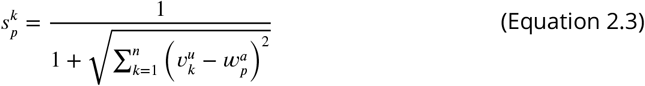

Unlike previous research investigating merging and fractionation in stroke survivors (Cheung et al., 2012; Hashiguchi et al., 2016; Pregnolato et al., 2025), we did not threshold the resulting values or define discrete events. Instead, we took the raw 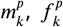 and 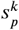 values as participant-specific measures of each physiological response at baseline and after treatment. Using these raw values, we applied a feature selection method (‘*f*_classif_’ function in the sklearn Python package) to identify statistically significant (i.e. *p* < 0.05) independent contributors to the corresponding patient clusters related to motor impairment (i.e. *R*_Pre_, *S*_Pre_, *R*_Post_, *S*_Post_). Finally, to provide a parsimonious model of these significant contributors to each partition, we included them as explanatory variables in a binary logistic regression model with forward selection (Wald’s criterion: inclusion < 0.05, exclusion > 0.1) (SPSS Statistics 28).

### Characterising rehabilitation effects on functional muscle interactions among responders and non-responders

To determine the effects of rehabilitation on functional muscle network structure among responders and non-responders, we firstly took the average functional interaction magnitude from the sparsified networks of participants affected and unaffected sides at each session. Using independent samples t-tests, we then determined statistical differences between patient groupings categorised based on the conventional MCID derived from the FMA-UE scale (i.e. 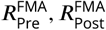 and 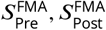), and on the patient response clustering’s established using our framework (i.e. *R*_(Pre-Post)_, *S*_(Pre-Post)_) (see ‘*Clustering stroke survivors based on impairment severity and therapeutic responsiveness*’ section of Materials and Methods). This enabled us to characterise the effects of rehabilitation on muscle network interdependencies and contrast between conventional and functional muscle network-based perspectives on therapeutic responsiveness.

Additionally, we aimed to quantify specific differences between responders and non-responders at each session at the level of individual muscle interactions. To do so, we employed a permutation-based approach to empirically derive interaction-specific significance thresholds and counteract family-wise error accumulation (Maris & Oostenveld, 2007). Briefly, for each grouping (i.e. clinical assessment-based and NIF) and muscle interaction across participants we randomly shuffled the group labels and determined statistical differences between the permuted groups using independent samples t-tests. This procedure was repeated *n* = 1000 times, generating a ground-truth distribution of test values from which the 95^th^ percentile value (i.e. *p* < 0.05) acted as the statistical significance threshold to compare against actual test values.

### Statistical analyses

The association between the extracted redundant- and synergistic pre- and post-session activation coefficients and both pre- and post-session FMA-UE scores respectively were assessed using univariate linear regression analyses (`“f_regression”` function in the `sklearn` python package). In this way, we could identify modular activation patterns that were independently associated with functional impairment. Additionally, we sought to determine if changes to the activation of specific redundant and synergistic components reflected differences between participants based on treatment group (i.e. PT vs. VR) and treatment responsiveness (i.e. MCID on the FMA-UE (> 5 points = responder, ≤ 5 points = non-responder) (Page et al., 2012). To do so, we firstly computed the Canberra distance (i.e. 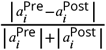) between corresponding pre- and post-session activation coefficients 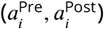 as a metric sensitive to small but potentially important proportional differences in modular activation across sessions. Using these distance values, we then determined differences between treatment responders and non-responders using independent t-tests. To control for the imbalanced groups, differences between experimental groups were examined instead using the Welch’s t-test. Significance was set for all statistical analyses here a priori to *p* < 0.05.

## Acknowledgments

David O’Reilly and Ioannis Delis were funded by the Biotechnology and Biological Sciences Research Council (BB/Y513799/1). This publication emanated from research supported by the European Union’s Horizon 2020 research and innovation programme (Marie Skłodowska-Curie grant agreement No 101034252).

## Data Availability

The data is available upon reasonable request.

## Supplementary Materials

### NIF Framework Application

The application of the NIF to the post-stroke cohort across both pre- and post-sessions identified five redundant (R1-R5 (Supplementary Materials Fig.1)) and seven synergistic (S1-S7 (Supplementary Materials Fig.2)) networks of co-occurring muscle interactions across affected and unaffected sides. We found a wide range of muscle interaction strengths (illustrated as the relative edge-widths on the human body models), principally contributing muscles (relative node sizes on human body model networks) and submodular affiliations (i.e. modules-within-modules) (node colours on the human body model networks) (see tables in Supplementary Materials Fig.1-2 for summary statistics including the range for the entire network (i.e. ‘*Full Network*’) and the ‘*Extraction of Redundant and Synergistic Muscle Networks*’ section of the Materials and Methods for further details). Interpreting these interaction strengths and principal muscles together with the submodular structure (see tables in Supplementary Materials Fig.1-2), we were able to attribute a biomechanical function to each extracted component.

Beginning with the redundant muscle networks (Supplementary Materials Fig.1), we found the biomechanical function of most components included the stabilisation of the shoulder girdle and/or elbow joint. These task-specific contributions to musculoskeletal stability were in some cases the predominant function of the muscle network (e.g. R3 (Affected-side), R5 (both sides)) and for others played a supportive role to elbow extension (e.g. R2 (both sides), R4 (Unaffected-side)) or forearm/shoulder rotation (e.g. R1 and R3 (both sides), R4 (Affected-side)). Examination of the submodular structure of the redundant muscle networks, which typically included 2-3 submodules, supported these functional interpretations. For example, for the affected-side R1 which related primarily to elbow extension and shoulder flexion (Fig.2), we found TM and TL were affiliated with separate submodules (green and red nodes respectively) including AD and LD (green nodes) and PD and BRACH (red nodes). This functional grouping reflects the distinct prime-mover and supportive contributions of the triceps brachii heads to elbow extension/shoulder flexion that are shared with specific elbow/shoulder muscles (i.e. TL contributes more to voluntary elbow extension than TM and is affiliated with shoulder/elbow stabilisers; TM contributes to shoulder stabilisation more so than TL and is coupled with shoulder flexors). Hence, knowing the activation state of TL or TM alone was sufficient to predict the motor task performed in an equivalent way as to observing the activation state of PD/BRACH or AD/LD respectively here. Differences were observed between the affected- and unaffected-sides of the redundant muscle networks indicative of motor impairment. R3 for instance (Supplementary Materials Fig.1), which involved interactions between SU and the shoulder musculature on both sides, displayed a specific reliance on PM on the affected-side whereas for the unaffected-side, the interactions were spread across PM, AD and LD, perhaps representing a loss of muscle selectivity at the shoulder level across the post-stroke cohort.

The extracted synergistic motor components (Supplementary Materials Fig.2) presented with widespread functional connections between proximal and distal musculature (e.g. S6 (Affected-side)). The extracted components were also less comparable across affected and unaffected sides compared to the redundant motor components (Supplementary Materials Fig.1), suggesting that the synergistic interactions more frequently reflected functional deficits induced by hemiparesis in the post-stroke cohort. Moreover, the submodular structure of the synergistic networks was more complex on the affected-side, ranging from 2-5 submodules compared to the 2-3 submodules typically found on the unaffected-side. The biomechanical function of several synergistic components was related to elbow flexion (i.e. S1, S3-S4 (both sides) and S2 (affected-side) (Supplementary Materials Fig.2)) in contrast to the elbow extensors common to the redundant components (Supplementary Materials Fig.1), while all extracted components on both sides were comprised of some form of task-specific joint stabilisation (i.e. shoulder girdle or elbow joint). This is intuitive as prime-mover and stabiliser muscles may also have synergistic roles (i.e. they predict complementary features of the motor task). The unaffected-side S7 exemplifies this intuition (Supplementary Materials Fig.2), comprising of the prime-mover AD during shoulder flexion coupled with several shoulder girdle stabilisers that offer complementary information regarding the specific orientation of the arm when executing the motor tasks. Similarly to the separate submodular affiliation of TM and TL in the affected-side R1 described previously (Supplementary Materials Fig.1), the affected-side S7 here comprised of separate affiliations for the biceps brachii heads (BS (orange nodes) and BL (black nodes) (Supplementary Materials Fig.2)). The complementary task information provided by BS here was more closely aligned with AD while BL was affiliated with BRACH, RM and Infra. These distinct functional affiliations were present despite a prominent functional interaction between BS and BL, suggesting that their complementary functions towards elbow flexion and shoulder girdle stabilisation to some extent overlapped.

### Comparison with conventional approaches

To more directly illustrate the advantages of the proposed framework, we carried out a standardised pre-processing of the EMG data in line with conventional muscle synergy analysis. This included rectification, low-pass filtration (cut-off: 20Hz) and smooth resampling of EMG waveforms to 50 timepoints. All data for each participant at each session was separately normalised by channel-wise variance, concatenated together and input into non-negative matrix factorisation (NMF) (*‘nnmf’* Matlab function, 10 replications) to extract 11 muscle synergies (*W*_1−11_ of Supplementary Materials Fig. 5(Left)) and their time-varying activations. The number of components to extract was determined in a conventional way as the number of components required to explain >75% of the data variance. The extracted muscle synergies included distinct shoulder- (e.g. *W*_2_), elbow (e.g. *W*_8_) and forearm-level (e.g. *W*_1_) muscle covariation patterns along with more isolated muscle contributions (e.g. *U T* in *W*_3_, *T L* in *W*_10_).

Regarding the clustering results of our framework and how they compare to conventional approaches, to facilitate this comparison we applied agglomerative clustering to the time-varying activation coefficients of all participants, trials, tasks separately for pre- and post-sessions and employed the *‘evalclusters’* Matlab function (Ward linkage clustering, Calinski Harabasz criterion, Klist search = 2:21) for each session. We identified two clusters both at pre-session (Criterion = 1.69) and post-session (Criterion = 1.81) as optimal fits to the population data (see Supplementary Materials Fig. 5(Right)). We found no associations between pre- or post-session cluster partitions and participants FMA-UE scores. Nevertheless, we did identify significant associations between the pre-session clusterings and *S*_*Pre*_ (*χ*^2^ = 7.08, *p* = 0.008) and between post-session clusterings and conventionally-defined treatment responders (*χ*^2^ = 4.2, *p* = 0.04). These findings, along with the similar two-way clustering structure found using the NIF, highlights important commonalities between these approaches.

To summarise the main advantages of our framework over this conventional approach:

•

- **Enhanced interpretability of extracted components and clusters**. As our framework maps muscle interactions to a specific task parameter, we yield population-level motor components that correspond more consistently to meaningful biomechanical and physiological functions that can be interpreted across the dimensions of the specified task parameter. The proposed clustering approach also offers enhanced interpretability, addressing key limitations in the application of clustering approaches to the activation space of conventional muscle synergy analysis (e.g. different activation timings) (Scano et al., 2017).
- **Integration of pairwise muscle relationships**. By incorporating muscle-pair level analysis, our framework captures coordinated interactions between primary and stabilising muscles—relationships that conventional NMF approaches overlook.
- **Separation of task-relevant and task-irrelevant activity**. The NIF isolates task-relevant coordination patterns, distinguishing them from task-irrelevant interactions driven by biomechanical or task constraints. On the other hand, task-relevant and -irrelevant muscle contributions are intermixed in conventional muscle synergy analysis.
- **Ability to identify complementary functional roles**. The NIF characterises whether muscle pairs act in similar or complementary ways, providing richer insight into motor control strategies.
- **Reduced dependence on variance-based optimisation**. Unlike conventional methods that rely on maximising variance explained, our framework allows detection of subtle but functionally significant interactions that contribute less to total variance.

## Supplementary Figures

**Figure 1.**
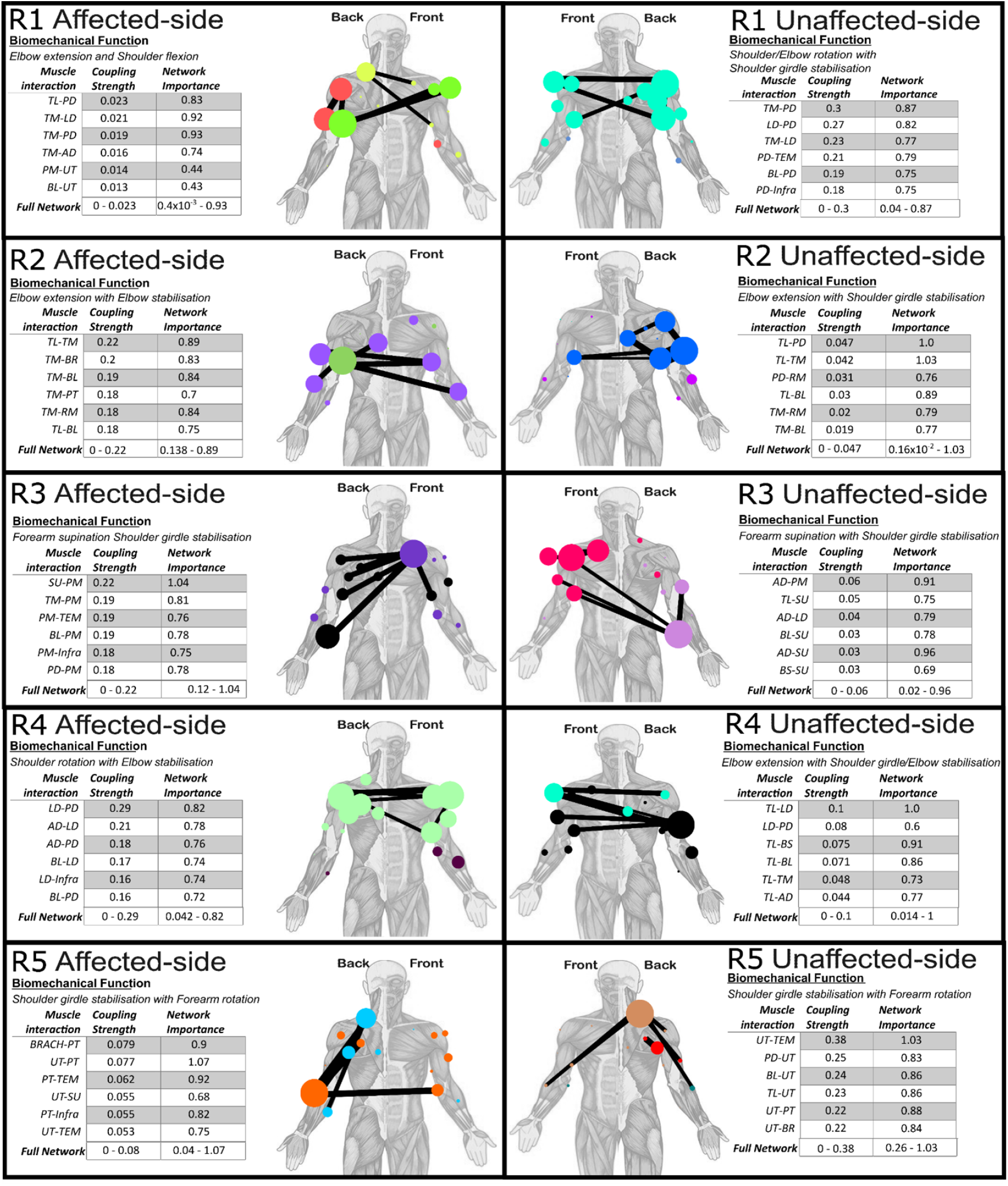
Five redundant network components (R1-R5) were extracted from the post-stroke cohort across pre- and post-sessions for both affected- and unaffected-sides. The 95th percentile of highest magnitude muscle interactions are illustrated for both sides on human body models along with the relative interaction strengths (edge-widths), relative network importances (node size) and sub-modular affiliations (node colour). The biomechanical function interpreted from each component accompanies each human body model and is supported by a table detailing the interaction strength of each muscle interaction along with their combined network importance (i.e. the sum of their eigenvector centrality).

**Figure 2.**
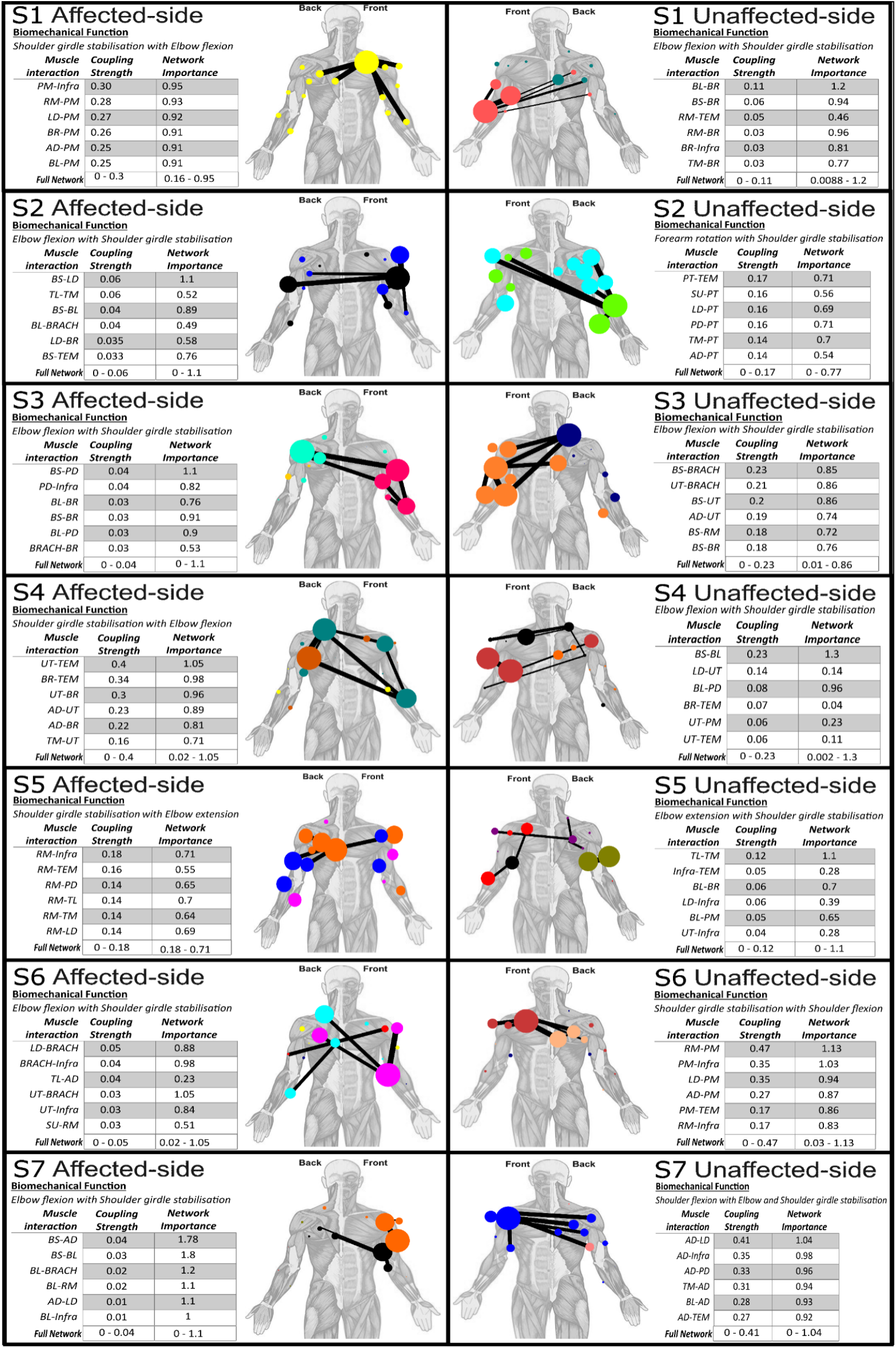
Seven synergistic network components (S1-S7) were extracted from the post-stroke cohort across pre- and post-sessions for both affected- and unaffected-sides. The 95th percentile of highest magnitude muscle interactions are illustrated for both sides on human body models along with the relative interaction strengths (edge-widths), relative network importances (node size) and submodular affiliations (node colour). The biomechanical function interpreted from each component accompanies each human body model and is supported by a table detailing the interaction strength of each muscle interaction along with their combined network importance (i.e. the sum of their eigenvector centrality).

**Figure 3.**
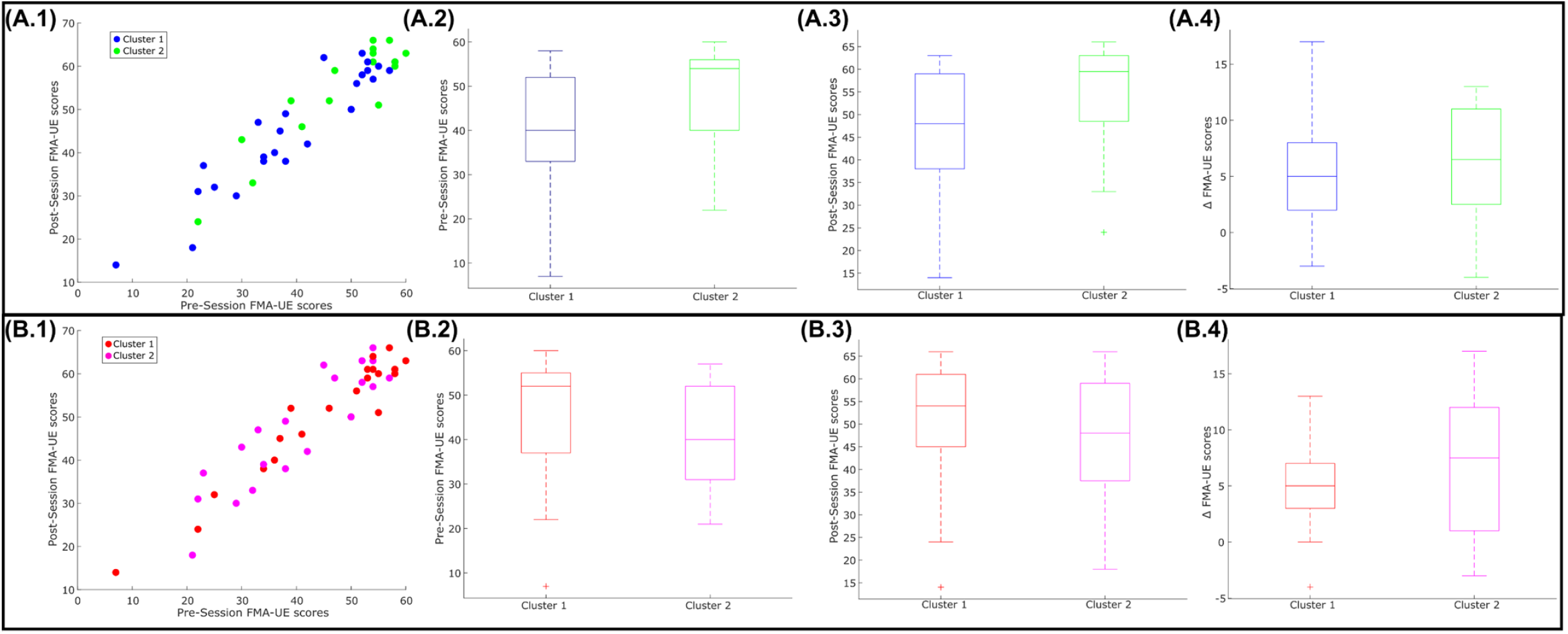
The dichotomous patient clusters derived from functionally-similar **(A)** and -complementary **(B)** as scatter plots with respect to post-session FMA-UE scores **(1)** and as box-plots with respect to pre-session FMA-UE **(2)**, post-session FMA-UE **(3)** and the change in FMA-UE scores (ΔFMA-UE) **(4)**. No significant differences could be found across **(A.2-4)** or **(B.2-4)** (p>0.05).

**Figure 4.**
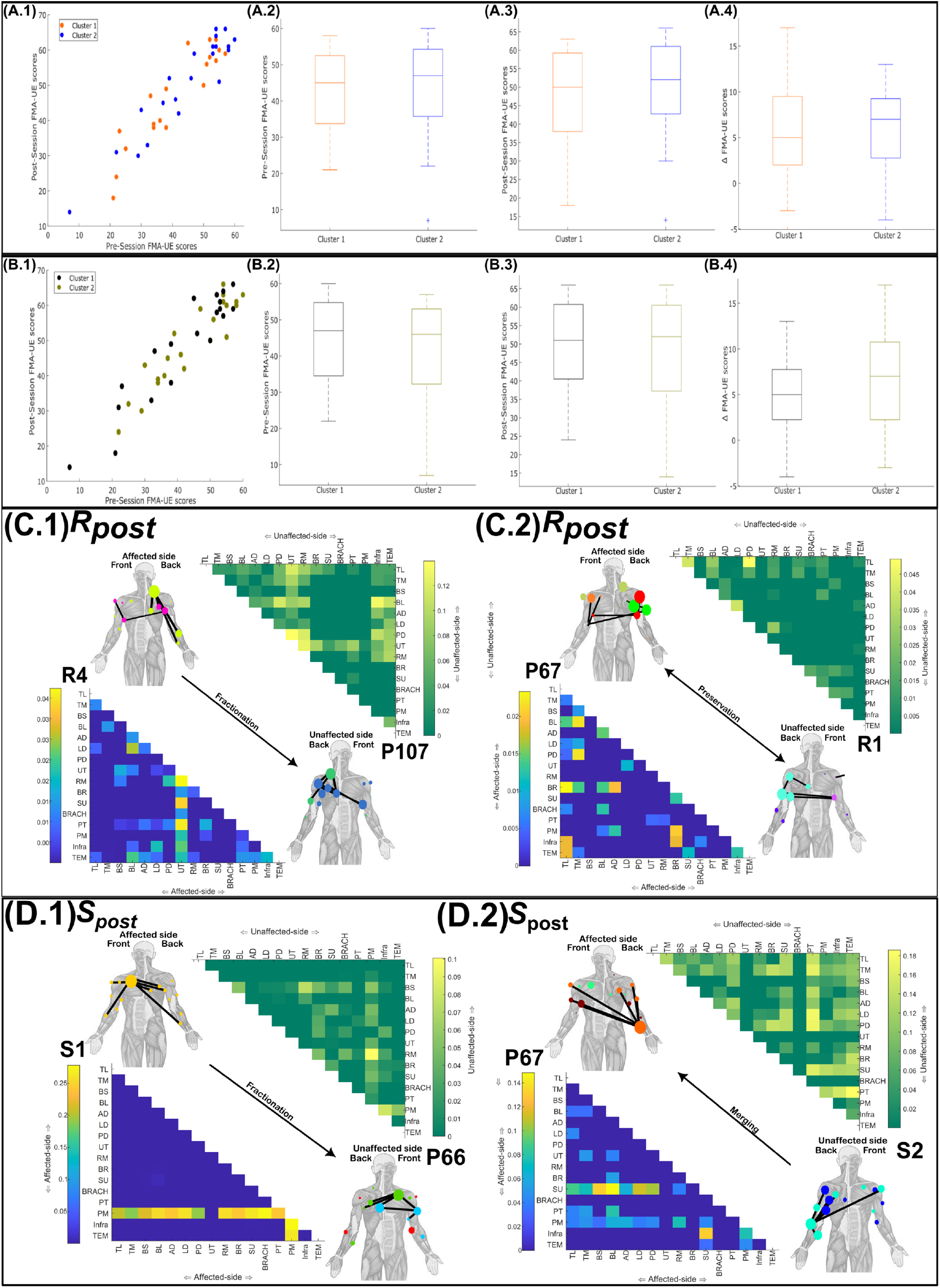
The identified patient clusters depicted with respect to post-session FMA-UE scores for (**A.1**) and (**C.1**). Boxplots illustrating the differences between the clusters identified in each partition for baseline FMA-UE scores (**A.2**-**B.2**), follow-up FMA-UE scores (**A.3**-**B.3**) and the change in FMA-UE scores from baseline to follow-up (i.e. ΔFMA-UE) (**A.4**-**B.4**). No significant differences between patient clusters were found across **(A-B.1-4)** (*p* > 0.05). The network components identified as significantly contributing to **(C.1-2)** and **(D.1-2)** through fractionation (affected-side (lower triangular matrix)), preservation and merging (both unaffected-side (upper triangular matrix)). For interpretation, a corresponding affected- or unaffected side network from a representative participant accompanies each significant network component (i.e. P66, P67 and P107). The submodular structure (node colour) and most proportionally significant (i.e. >95^th^ percentile) muscle couplings (network edges) are also illustrated for each network on human body models. **(C.1)** Fractionation of R4 and **(C.2)** preservation of R1 explained participants’ affiliation with cluster 1 of (*β* = -2.32±0.99 (*p* = 0.019), *β* = -15.2±6.5 (*p* = 0.018) respectively), classifying 81% of participants correctly. **(D.1)** Fractionation of S1 (*β* = 26.7±12.1 (*p* = 0.027)) and **(D.2)** merging of S2 (*β* =27.3±11.8 (*p* = 0.021)) explained participants affiliation with cluster 2, together classifying 76.2% of participants correctly.

**Figure 5.**
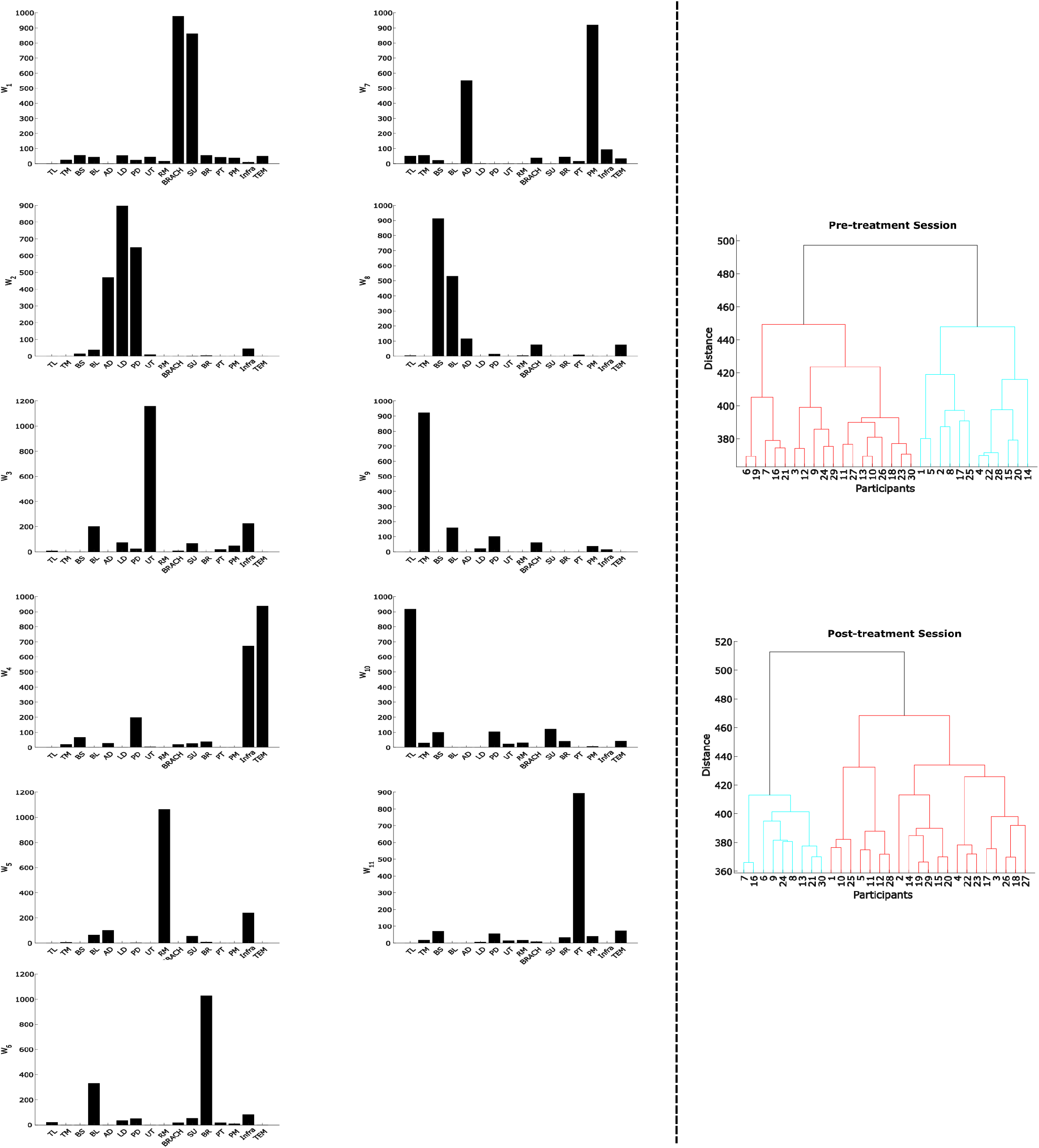
To perform a direct comparison with existing approaches, we applied the conventional muscle synergy analysis approach based solely on non-negative matrix factorisation (NMF) to the post-stroke cohort. **Left**: We extracted 11 muscle synergies (*W*_1−11_) and performed agglomerative clustering on the time-varying activation coefficients for pre- and post-sessions separately. **Right**: Statistical analysis of the clusters identified revealed no significant differences for pre- or post-session FMA-UE scores.

